# Reconstitution of Kinesin-1 Activation

**DOI:** 10.1101/2021.03.12.434960

**Authors:** Kyoko Chiba, Kassandra M. Ori-McKenney, Shinsuke Niwa, Richard J. McKenney

## Abstract

Autoinhibition is an important regulatory mechanism for cytoskeletal motor proteins. Kinesin-1 (kinesin hereafter), the ubiquitous plus-end directed microtubule motor, is thought to be controlled by a complicated autoinihibition mechanism, but the molecular details remain unclear. Conformational changes mediated by intramolecular interactions between the C-terminal tail and N-terminal motor domains of the kinesin heavy chain (KHC) are proposed to be one facet of motor regulation. The dimeric KHC also binds two copies of the kinesin light chains (KLCs), which have been implicated in both autoinhibition and cargo-dependent activation of the tetrameric motor complex, although the precise mechanisms remain opaque. Using in vitro reconstitution, we show that the KLC strongly inhibits the kinesin-microtubule interaction via an independent mechanism from the tail-motor interaction within KHC. Kinesin cargo-adaptor proteins that bind KLC activated processive movement of the kinesin tetramer but the landing rate of these activated complexes remained low. The addition of MAP7, which specifically binds to the KHC, strongly enhanced activated motor motility by dramatically increasing the landing rate and processivity of the activated kinesin motors. Our results support a model whereby the activity of the kinesin tetramer is regulated by independent tail- and KLC-based inhibition mechanisms, and that cargo-adaptor binding to the KLC directly releases both of these inhibitions. However, we find that a third component, a non-motor MAP is required for robust activity of the activated motor. Thus, human kinesin activity is regulated by a two-factor mechanism comprised of intramolecular allosteric regulation, as well as intermolecular kinesin-adaptor and kinesin-MAP interactions.

## Introduction

Microtubule motor proteins from the kinesin and dynein families play important roles in the intracellular transport of proteins, ribonucleoproteins, membranous organelles, and many other types of cargos (Hirokawa et al., 2009; Vale, 2003). Autoinhibition is a common mechanism to restrict the activity of the motors in the absence of their cargoes so that cells avoid useless expenditure of ATP and aberrant movement of cellular cargos. Recent in vitro reconstitution and structural biology studies have revealed a wealth of knowledge about how cytoplasmic dynein, the only minus-end directed transport motor in animals, assumes an autoinhibited conformation that is relieved by cargo adapter molecules and a ubiquitous cofactor, dynactin (McKenney et al., 2014; Reck-Peterson et al., 2018). On the other hand, while kinesin is the most ubiquitous plus-end directed motor in cells, we currently lack such a detailed understanding of how this motor is regulated. While prior studies have found that the kinesin motor is also regulated by autoinhibition (Dietrich et al., 2008; Kaan et al., 2011; Verhey et al., 1998), our understanding of the mechanisms underlying kinesin inhibition and activation now dramatically lags behind our understanding of dynein regulation.

Kinesin predominantly exists as heterotetrameric complex composed of a dimer of kinesin heavy chains (KHC) bound to two copies of the kinesin light chain (KLC) subunit (Hirokawa et al., 1989; Vale et al., 1985). The N-terminal motor domain of kinesin uses the energy of ATP hydrolysis to produce force along microtubules. The motor domain (or head) is followed by a stalk, which is a series of coiled-coil domains separated by predicted flexible breaks, and by a C-terminal tail domain which is predicted to be largely disordered (Seeger et al., 2012). The C-terminal region of the stalk domain provides a binding site for the KLC (Diefenbach et al., 1998; Verhey et al., 1998), The KLC binds to the KHC via its N-terminal predicted coiled-coil (Verhey et al., 1998) and links various cargoes to kinesin through its C-terminal TPR domain (Pernigo et al., 2013). A pool of kinesin exists as KHC dimers and presumably functions in the absence of the KLC (DeLuca et al., 2001; Gyoeva et al., 2004; Palacios and St Johnston, 2002). However, the number of cellular cargos proposed to be transported by KHC dimers, versus kinesin tetramers, remains relatively low (Glater et al., 2006; Palacios and St Johnston, 2002; van Spronsen et al., 2013), indicating the kinesin tetramer is the predominant plus-end directed cargo-transport complex in cells.

In humans, three KHC (KIF5A-C) and four KLC (KLC1-4) genes comprise differentially expressed, biochemically distinct, isotypes of kinesin tetramers. The isotypes show distinct tissue expression patterns and produce unique phenotypes in null mutant mice. KIF5B and KLC2 are ubiquitously expressed whereas KIF5A, KIF5C and KLC1 are neuronally enriched (Kanai et al., 2000; Rahman et al., 1998). KLC3 is expressed in testis and little is known about KLC4 (Junco et al., 2001). Mice lacking either KIF5A or KIF5B are lethal, but KIF5C null mice are viable (Tanaka et al., 1998) (Kanai et al., 2000; Xia et al., 2003). In humans, mutations in KIF5C result in neurodevelopmental pathologies (Poirier et al., 2013; Willemsen et al., 2014). Additionally, mutations in KIF5A cause various neurodegenerative diseases (e.g. SPG10, Charcot-Marie-Tooth disease 2, amyotrophic lateral sclerosis (ALS) (Crimella et al., 2012; Nicolas et al., 2018; Reid et al., 2002). While these observations suggest functional differences exist among KIF5 isotypes, previous studies have analyzed different KHCs from different species and the relative biochemical and biophysical properties of different kinesin isotypes have not been systematically compared.

Several studies have proposed a model by which the motility of the KHC is regulated by a direct interaction between the tail domain and the motor domain (Coy et al., 1999; Friedman and Vale, 1999; Kaan et al., 2011; Stock et al., 1999). It postulates that the kinesin molecule flexes at a “hinge” in the middle of the coiled-coil stalk region, enabling a short, charged peptide within the tail, called the isoleucine–alanine–lysine (IAK) motif, to associate with the motor domain and inhibit its enzymatic and microtubule-binding activities. A single IAK motif added in trans was found to cross-link the two motor domains within a kinesin dimer crystal structure (Hackney and Stock, 2000; Kaan et al., 2011). This interaction is believed to restrict the relative movement of the motor domains, preventing movement along microtubules (Cai et al., 2007).

The KLC is also involved in autoinhibition of kinesin. The presence of KLC strongly decreases microtubule-binding and motility of the KHC in both biochemical and cellular assays (Verhey et al., 1998). It remains unclear whether KLC strengthens the tail-inhibition or independently regulates KHC activity. FRET experiments implied that the presence of the KLC keeps kinesin in the proposed folded state, but also forces the motor domains apart (Cai et al., 2007). This function of KLC partially disagrees with the tail-inhibition model, since binding of the IAK domain locks the motor domains in close proximity to each other (Kaan et al., 2011). The autoinhibited kinesin is proposed to be activated by the binding of several distinct adaptor proteins (Blasius et al., 2007; Fu and Holzbaur, 2013; Sun et al., 2011; Twelvetrees et al., 2019; Twelvetrees et al., 2016; Verhey et al., 2001) to either the KHC, KLC, or both. However, these experiments were performed in cell lysates with overexpressed proteins of interest, and thus it remains unclear if adaptor proteins directly activate kinesin in the absence of other cellular factors.

A series of biochemical and structural studies have uncovered two consensus KLC binding sequences found within numerous cellular proteins that are thought to act as cargo-adapter molecules for kinesin (Cross and Dodding, 2019; Pernigo et al., 2013). The isolated C-terminal TPR domains of KLC bind to either W-acidic or Y-acidic motifs with relatively low affinity (~5 μM and 0.5 μM respectively, (Pernigo et al., 2018; Pernigo et al., 2013; Yip et al., 2016)), suggesting that the formation of kinesin-cargo complexes is also regulated by autoinhibition. Despite the low affinity *in vitro*, W-acidic peptides, or small molecule mimetic drugs (Randall et al., 2017), appear sufficient to drive kinesin activity within living cells, suggesting that cargo-adapter binding to the KLC could directly activate the autoinhibited motor. However, Y-acidic peptides do not show such activity on their own, but instead may require the co-binding of other molecules to activate kinesin in cells or cell lysates (Blasius et al., 2007; Kawano et al., 2012). Despite these findings, direct evidence that binding of W-acidic or Y-acidic peptides leads to motor activation is lacking.

In this paper, we systematically study the autoinhibition of dimeric and tetrameric kinesin motor complexes from all human KHC isotypes using biochemical reconstitution and single molecule assays. We confirm the IAK domain of KHC strongly represses the microtubule binding of KHC, although the strength of the autoinhibition varies by KHC isotype. Tetramer formation with the KLC more strongly attenuates the kinesin-microtubule interaction, in agreement with previous results (Verhey et al., 1998). Interestingly, widely utilized mutants that are thought to abolish the tail-inhibition mechanism within the KHC, are still strongly repressed by KLC. This suggests that the mechanism of KLC-inhibition is independent of the reported KHC tail-inhibition mechanism.

We further show that the autoinhibited kinesin-1 is partially activated by purified nesprin-4, a cargo-adapter that contains a W-acidic motif. However, nesprin-4 binding only mildly stimulates the motor’s landing rate onto microtubules. The microtubule association of the activated motor complex is greatly enhanced by the inclusion of the kinesin-interacting microtubule-associated protein (MAP), MAP7 (Monroy et al., 2018; Sung et al., 2008). Thus, our results reveal that neither cargoadapter binding, nor MAP7 recruitment is sufficient to fully activate the kinesin motor. Instead, a two-factor activation mechanism composed of adapter-binding coupled with specific recruitment by a MAP is utilized by complex eukaryotic kinesin motors. These results have strong parallels with the dynein activation mechanism (McKenney et al., 2014; McKenney et al., 2016; Schlager et al., 2014), suggesting that cargo-adapter mediated release of autoinhibition coupled with MAP recruitment to microtubules, is a generalized mechanism favored by evolution for the regulation of long-distance transport by mammalian microtubule motor proteins.

## Results

### Direct Comparison of Human KIF5 Isotypes

While the biophysical parameters and structural mechanism of KIF5 movement has been studied in great detail (Cross, 2016; Gennerich and Vale, 2009; Hancock, 2016), these studies have utilized diverse species, kinesin isotypes, and assay conditions, complicating a direct comparison between the three distinct human KHC isotypes. Therefore, we first set out to compare the motor activity of all three human isotypes of the KHC (KIF5). We utilized tail-truncated, but natively dimerized and constitutively active motor constructs based on the bacterially expressed KIF5B K420 construct (Shimizu et al., 2000)(Fig. 1A, B). Because this truncation boundary contains only the minimum amount of the stalk domain required for dimerization, these constructs should report directly on the biophysical properties of the unregulated kinesin motor domain. All constructs contained a C-terminal mScarlet-StrepII cassette for purification and observation in single molecule assays.

**Figure 1.**
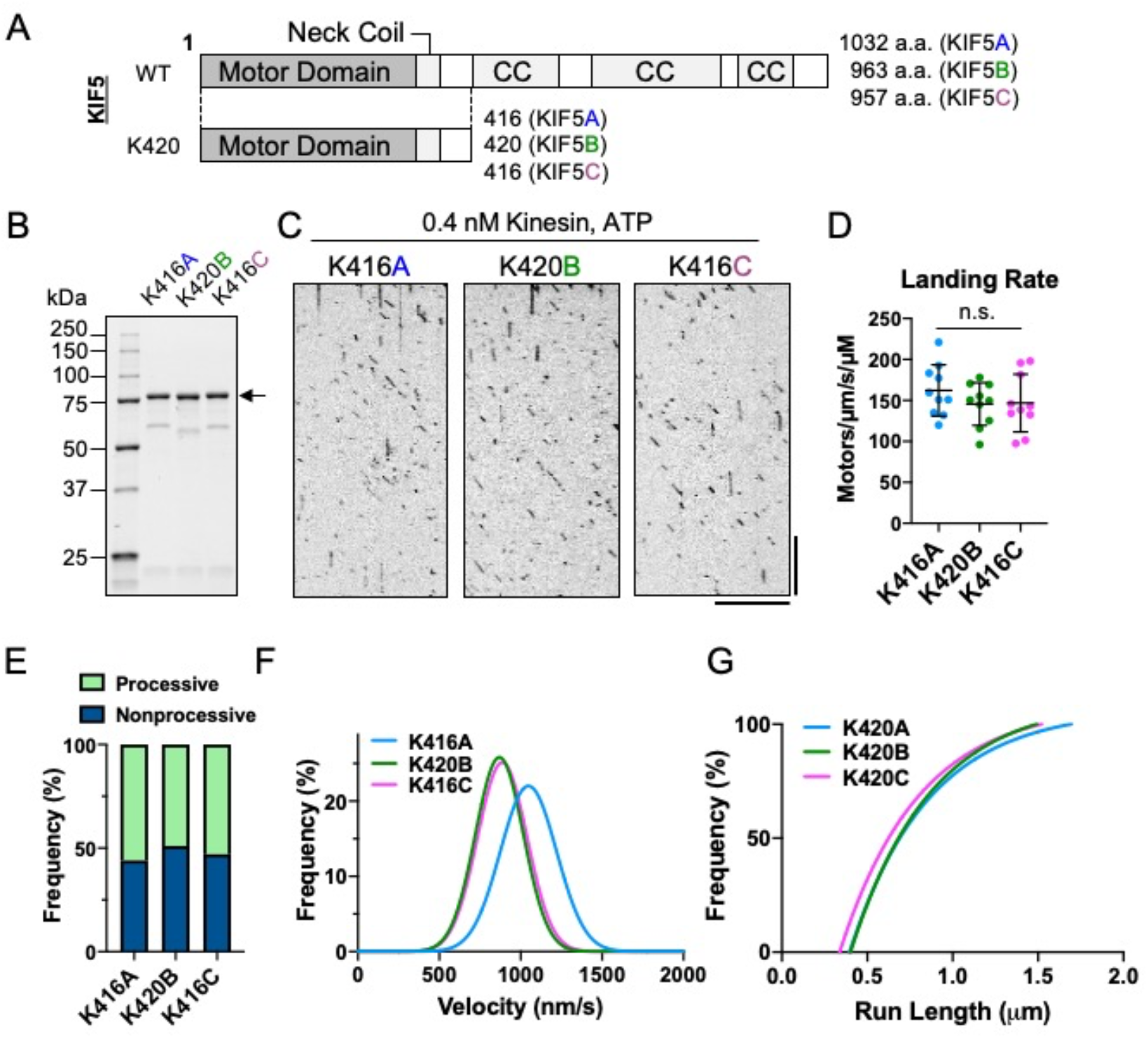
Characterization of truncated, constitutively active human kinesin-1 isotypes. **(A)** Schematic of full-length KIF5 and K420 constructs. Major domains are indicated. CC: coiled-coil. **(B)** Coomassie blue stained gel showing purified K420 proteins. Arrow indicates the full-length K420 proteins (74, 75 and 74 kDa for K420A, K420B and K420C, respectively). **(C)** Kymographs of mScarlet-tagged K420 motors moving along microtubules. Scale bars: 10 s (vertical) and 10 μm (horizontal). **(D)** Measured landing rates of each motor. Lines: mean ± SD (162 ± 31, 146 ± 26 and 147 ± 35 /μm/s/μM for K420A, K420B and K420C, respectively). One-way ANOVA followed by Tukey’s multiple comparison test was performed. n.s., not significant. N = 2 from two independent preparations. n = 10 microtubules measured per chamber. **(E)** Measured frequency of processive and non-processive (static or diffusive) events. Processive runs were 56% (K420A), 49% (K420B) and 53% (K420C), respectively. n = 715, 573 and 679 molecules. **(F)** Gaussian fits of motor velocities. Mean ± SD: 1048 ± 173 nm/s (K420A), 870 ± 147 nm/s (K420B) and 887 ± 152 nm/s (K420C). n = 397, 282 and 359 molecules, respectively. **(G)** One-phase exponential decay fits based on 1-cumulative frequency distribution of run lengths. Decay constant (τ, run length) were 0.46 μm for K420A, 0.47 μm for K420B and 0.46 μm for K420C.

The motor domains of KIF5A, B, and C are overall more than 70% identical to one another, suggesting the core motor mechanism is highly conserved between human isotypes. Indeed, we observed highly similar behavior of all three isotypes in our single molecule assay where motors landed on microtubules and often moved processively along them (Fig. 1C-G). Importantly, the landing rate of the truncated motors was very high and similar between all isotypes (Fig. 1D). Additionally, a similar fraction of motors that landed on the microtubule transitioned to processive movement for all isotypes (56%, 49% and 53% for K416A, K420B and K416C, Fig. 1E). Interestingly, the velocity of the KIF5A construct (1048 ± 173 nm/sec., s.d) was significantly faster than that of KIF5B or KIF5C (870 ± 147 and 887 ± 152 nm/sec., s.d. respectively. Fig. 1F), implying the mechanochemistry of KIF5A is unique among human KIF5 isotypes. The fitted run lengths of all three isotypes were also highly similar (0.46, 0.47 and 0.46 μm for KIF5A, B, C respectively. Fig. 1G). Thus, under identical assay conditions, the three human KIF5 motors have largely similar biophysical properties, but KIF5A displays a uniquely faster mechanochemistry, which may conceivably be related to its specialized cellular functions exclusively within neurons (Niclas et al., 1994).

### Isotype Specific Autoinhibition of Kinesin Dimers

Because the largest difference in primary sequence between human KIF5 isotypes lies within the distal C-terminal tail domains (< 20% identity in the tail region, Fig. 2A), we generated and purified recombinant full-length KHC isotypes using the baculovirus expression system (Fig. 2B). All three human isotypes were tagged with a C-terminal mScarlet-strepII tag for purification and visualization. We first characterized the oligomeric state of the purified proteins in solution using size-exclusion coupled with multi-angled light scattering (SEC-MALs). We found that human KIF5B and KIF5C isotypes existed predominantly as dimeric complexes, as expected, because their measured molecular mass was within ~ 20% of the predicted dimeric value for each (Fig. 2C-D). However, KIF5A deviated from its predicted dimeric molecular mass by greater than 20%, suggesting that a significant proportion of KIF5A molecules may exist in a more complex oligomeric state (Fig. 2C-D).

**Figure 2.**
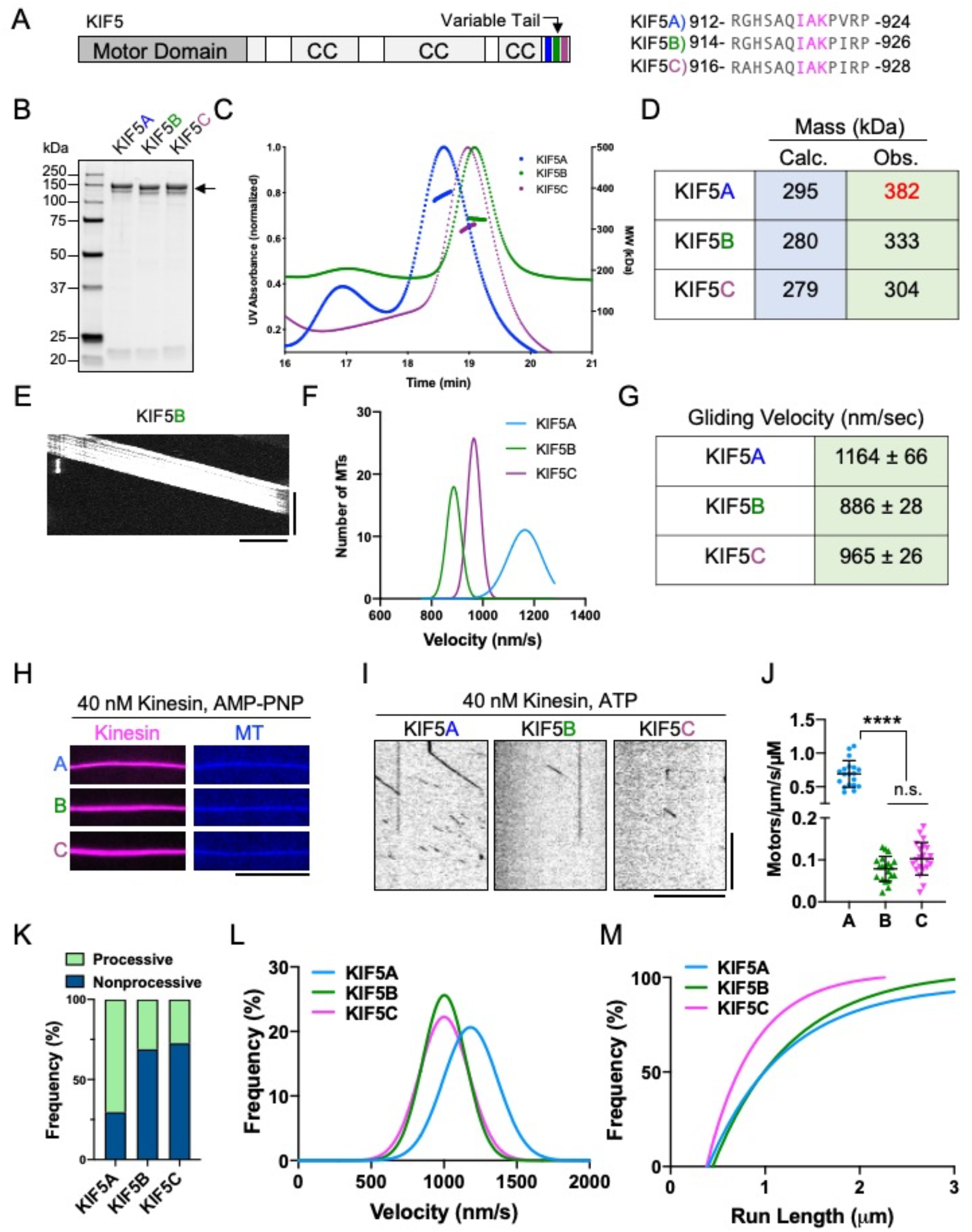
Charcaterization of full-length human KIF5 isotypes. **(A)** Schematic of full-length KHC and alignment of the conserved IAK motif (magenta) and its surrounding residues of KIF5A-C are shown. CC: coiled-coil. **(B)** Coomassie blue stained gel of purified KHC proteins. Arrow indicates the full-length KHC proteins (147, 140 and 140 kDa for KIF5A, KIF5B and KIF5C, respectively). **(C)** Chromatogram and MALS fitting of KIF5A (blue), KIF5B (green) and KIF5C (purple). Normalized UV absorbance at 280 nm (dotted lines) and molar masses are plotted. **(D)** Table summarizing the calculated theoretical and the experimentally observed masses for each motor. **(E)** Kymograph showing smooth microtubule movement in the gliding assay with the KIF5B motor. Scale bars are 20 s (vertical) and 5 μm (horizontal). **(F)** Gaussian fits of the velocity histograms for KIF5A (blue), KIF5B (green) and KIF5C (purple) from the gliding assay. Mean ± SD: 1160 ± 66 nm/s (KIF5A), 887 ± 26 nm/s (KIF5B) and 965 ± 27 nm/s (KIF5C). n = 90, 70 and 90 microtubules (MTs), respectively. **(G)** Table summarizing the average gliding velocities of KIF5A-C. **(H)** TIRF images of KIF5A-C motors (magenta) bound to microtubules (blue) in the presence of 2 mM AMP-PNP. Scale bar = 10 μm. **(I)** Kymographs of single KIF5A-C motors moving along microtubules in the presence of 2 mM ATP. Scale bars are 10 s (vertical) and 10 μm (horizontal). **(J)** Landing rates of KIF5A-C. Lines: mean ± SD (0.69 ± 0.20, 0.08 ± 0.03 and 0.10 ± 0.04 /μm/s/μM for KIF5A, KIF5B and KIF5C, respectively). n = 19, 20 and 21 MTs. N = 2 experimental replicates from two independent preparations. One-way ANOVA followed by Tukey’s multiple comparison test was performed. **** = P < 0.0001; n.s., not significant. **(K)** Frequency of processive non-processive (static or diffusive) events observed. Processive runs for KIF5A, KIF5B and KIF5C were 70%, 31% and 27%, respectively. n = 1165, 1209 and 901 molecules. **(L)** Gaussian fits of KIF5 velocities. Mean ± SD: 1182 ± 187 nm/s (KIF5A), 1002 ± 149 nm/s (KIF5B) and 1001 ± 170 nm/s (KIF5C). n = 820, 376 and 247 molecules, respectively. **(M)** One-phase exponential decay fits based on 1-cumulative frequency distribution of run lengths. Decay constants (τ, run length) were 0.82 μm for KIF5A, 0.82 μm for KIF5B and 0.51 μm for KIF5C.

We then tested the motor activities of our kinesin preparations using a multimotor microtubule-gliding assay in which the motors are attached nonspecifically to a glass surface. For all isotypes, we observed smooth, continuous movement of microtubules across the glass surface (Fig. 2E), confirming that all three recombinant motors were active. The distribution of microtubule gliding velocities were very similar for KIF5B and KIF5C, although KIF5C was slightly faster (887 ± 26 and 965 ± 27 nm/s, s.d. for KIF5B and KIF5C respectively. Fig. 2F-G). However, we again observed a significantly faster velocity for KIF5A (1160 ± 66, s.d.), compared to the other isotypes (Fig. 2F-G), further suggesting the mechanochemistry of KIF5A is unique. These results show that recombinant human kinesin dimers are highly active motors with largely overlapping biochemical and biophysical parameters, although KIF5A’s mechanochemistry is unique among human KIF5 isotypes.

Next, we studied the single molecule behavior of human KHC isotypes using total internal reflection fluorescence (TIRF) microscopy. All three human KHC isotypes contain the highly conserved ‘IAK’ motif located within their tail domains (Fig. 2A), which plays a direct role in an intramolecular interaction between the tail domain and the motor domains (Dietrich et al., 2008; Kaan et al., 2011). Based on these prior findings, we expected the microtubule-binding and motor activities of the full-length motors to be strongly attenuated as compared to the tail-truncated constructs (Fig. 1). We first confirmed that all three isotypes bind strongly to microtubules in the presence of the ATP analogue, 5’-adenylylimidodiphosphate (AMP-PNP, Fig. 1H). Thus, the presence of the intact tail domain does not preclude the binding of the purified motors to microtubules, in agreement with prior findings in cell lysates (Vale et al., 1985; Verhey et al., 1998). In the presence of ATP, we observed very few binding events along microtubules, even at concentrations 100-fold higher than those used for the tail-truncated constructs (Fig. 2I, compare to Fig. 1C). Quantification of the microtubule-landing rate revealed that once again, KIF5A displayed unique behavior, having a seven to ten-fold higher landing rate than KIF5B and KIF5C, respectively (Fig. 2J), and a much higher fraction of processive motors (Fig. 2K). However, the landing rates of the full-length motors was several hundred-fold lower than those for the tail-truncated constructs (Fig. 1D), confirming that the presence of the tail domain strongly impacts the microtubule association rate of the motor.

Close inspection of the kymographs revealed that motile molecules of KIF5A were sometimes brighter than those of the other isotypes (Fig. 2I), consistent with the SEC-MALs findings that our preparation of KIF5A contains a fraction of higher-order oligomers (Fig. 2C-D). Quantification revealed the brighter KIF5A molecules were 2 ± 0.65 (s.d) −fold brighter (n = 206 molecules from N = 2 trials) than the dimmer KIF5A molecules in the same chamber. We conclude that the brighter population of KIF5A molecules may represent tetrameric complexes, in support of our SEC-MALs data (Fig. 2C, D). Consistently, the brighter KIF5A molecules often traveled longer distances than the dimmer molecules within the KIF5A population or compared to KIF5B or KIF5C (Fig. 2I, L). Indeed, quantification of velocity (1182 ± 187, 1002 ± 149, 1001 ± 170 nm/s, s.d. for KIF5A, B, and C respectively) further revealed distinctly fast behavior for KIF5A (Fig. 2L). We note that the single molecule velocities are similar to the velocities of microtubule gliding in the multi-motor assay, suggesting that the microtubule-stimulated enzymatic rate of kinesin is not strongly affected by the presence of the tail domain. However, quantification of motor run-lengths (0.82, 0.82 and 0.51 μm for KIF5A, B, and C respectively) revealed distinctly shorter excursions for KIF5C (Fig. 2M). These results confirm that the tail domain of kinesin largely inhibits its processive movement, consistent with prior findings (Friedman and Vale, 1999). However, we extend these previous findings by revealing that the efficiency of tail-mediated inhibition varies between human KHC isotypes. Therefore, the unique tail sequences of each KHC contribute to the overall level of basal activation of the motor. Our results also suggest that the KIF5A dimer is predisposed to form higher-order oligomers, in our conditions, which we do not observe for KIF5B or KIF5C.

### Association with the KLC Further Inhibits the Kinesin-Microtubule Interaction

Most kinesin molecules in the cell are thought to exist as a tetrameric complex with the KLC and evidence for a role of KHC dimers in cargo transport is limited. Previous studies have suggested that binding of the KLC to the KHC strongly represses the kinesin-microtubule association (Verhey et al., 1998) and enzymatic activity (Coy et al., 1999; Stock et al., 1999). We prepared recombinant kinesin tetramers with all three human KHC isotypes co-expressed with the brain-specific KLC isotype, KLC1 (Kawano et al., 2012; Verhey et al., 1998), using a multi-gene baculovirus (Fig. 3A-B). We measured a stoichiometric ratio of ~ 1.0:0.8 for KHC:KLC1 by coomassie blue staining, suggesting that the majority of the KHC was bound to KLC1. SEC-MALs analysis of the purified complexes further revealed a homogenous population of tetrameric motor complexes for all three KHC isotypes, which measured within 10% of the expected molecular weight (Fig. 3C-D). Thus, association with the KLC stabilizes the dimeric form of KIF5A, which tends to form higher-order oligomers in its absence (Fig. 2C-D).

**Figure 3.**
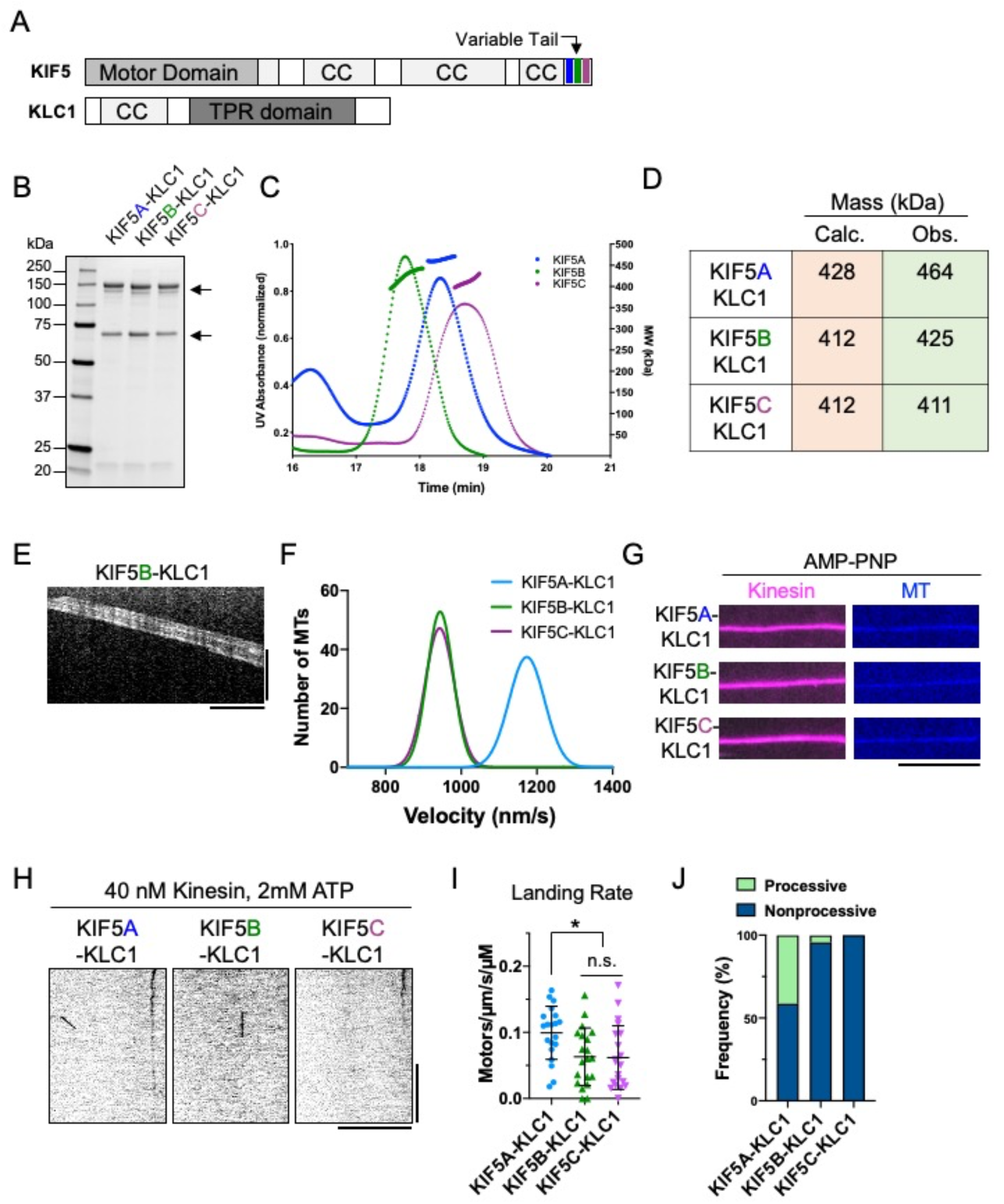
Characterization of the KIF5-KLC tetramer. **(A)** Schematic of full-length KHC and KLC1. CC: coiled-coil **(B)** Coomassie blue stained gel showing purified tetramers. Arrows indicate the full-length proteins (147, 140 and 140 kDa for KIF5A, KIF5B and KIF5C. 66 kDa for KLC1). **(C)** Chromatogram and MALS fitting (size exclusion chromatography coupled to multiangle light scattering) of KIF5A-KLC1 (blue), KIF5B-KLC1 (green) and KIF5C-KLC1 (purple). Normalized UV absorbance at 280 nm (dotted lines) and molar masses are plotted. **(D)** Table summarizing the calculated theoretical and the experimentally observed masses for each motor. **(E)** A kymograph of MT gliding assay using KIF5B-KLC1 tetramer. Scale bars are 20 s (vertical) and 5 μm (horizontal). **(F)** Gaussian fits of the velocity histograms for KIF5A-KLC1 (blue), KIF5B-KLC1 (green) and KIF5C-KLC1 (purple) from the microtubule gliding assay. Mean ± SD: 1176 ± 47 nm/s (KIF5A-KLC1), 943 ± 33 nm/s (KIF5B-KLC1) and 944 ± 36 nm/s (KIF5C-KLC1). n = 90 MTs, respectively. **(G)** TIRF images of KIF5-KLC1 tetramers (magenta) bound to microtubules (Blue) in the presence of 2 mM AMPPNP. Scale bar is 10 μm. **(H)** Kymographs of KIF5-KLC1 complexes on microtubules in the presence of 2 mM ATP. Scale bars are 10 s (vertical) and 10 μm (horizontal). **(I)** Measured landing rates of KIF5-KLC1 tetramers. Lines: mean ± SD (0.10 ± 0.04, 0.06 ± 0.04 and 0.06 ± 0.05 /μm/s/μM for KIF5A-KLC1, KIF5B-KLC1 and KIF5C-KLC1). n = 20 MTs, respectively. N = 2 experimental replicates from two independent preparations. One-way ANOVA followed by Tukey’s multiple comparison test was performed. * Adjusted P < 0.05; n.s., not significant. **(J)** Measured frequency of processive runs and non-processive (static or diffusive) movement. Processive runs were 41%, 4% and 0% for KIF5A-KLC1, KIF5B-KLC1 and KIF5C-KLC1, respectively. n = 290, 91 and 89 molecules, respectively.

We then tested the activity of the recombinant kinesin tetramers in the multimotor microtubule gliding assay. Similar to the KHC dimers (Fig. 2E), all three isotypes powered the smooth continuous gliding of microtubules when directly attached to a glass surface (Fig. 3E). The gliding velocities of tetramers containing KIF5B or KIF5C isotypes were nearly indistinguishable (Fig. 3F), and very similar to the gliding velocities of KHC dimers containing these isotypes (Fig. 2G). However, we once again observed a significantly faster velocity for tetramers containing the KIF5A isotype (Fig. 3F), further confirming the unique mechanochemistry of this isotype. The highly similar gliding velocities for dimeric and tetrameric kinesin complexes is consistent with a previous report for isolated kinesin dimers and tetramers from brain (Hackney, 1991). Thus, the KLC subunit stabilizes the dimeric form of KIF5A, but does not impact the mechanochemistry of the KHC in a multi-motor assay, in which the motors may be forced into an active conformation by binding to the glass surface.

Next, we assessed the ability of the purified kinesin tetramers to bind to microtubules in the presence of AMP-PNP. As expected (Vale et al., 1985; Verhey et al., 1998), the presence of the KLC did not inhibit the binding of the purified motors to microtubules (Fig. 3G). In the presence of ATP, we observed very little interaction of the purified tetramers with microtubules at concentrations 100-fold higher than those used for the truncated motors (Fig. 3H, compare to Fig. 1C). Indeed, the quantified landing rates of all three tetrameric motors (0.10 ± 0.04, 0.06 ± 0.04 and 0.06 ± 0.05 motors nm^-1^ min^-1^, s.d. for KIF5A-KLC1, KIF5B-KLC1 and KIF5C-KLC1 respectively, Fig. 3I) was several hundred-fold lower than for tail-truncated motors (Fig. 1D), and comparable to the landing rates of the kinesin dimers (Fig. 2J). However, we found that the KIF5A-KLC1 tetramer had a seven-fold lower landing rate than the KIF5A dimer, revealing that KLC-induced stabilization of the KIF5A dimer leads to a more repressed motor with similar microtubule-association kinetics to KIF5B and KIF5C dimers or tetramers.

Although microtubule-association events were rare, we quantified the percentage of events that were processive versus static or diffusive and again observed that KIF5A tetramers showed a much larger fraction of processive motors compared to KIF5B or KIF5C tetramers (Fig. 3J). Thus, in agreement with prior studies in cell lysates (Verhey et al., 1998), association of the KHC with the KLC leads to largely repressed tetrameric motor complex regardless of the KHC isotype. Given the higher ratio of KIF5A-KLC1 complexes that moved processively, we were able to quantify enough processive events to derive velocity and run-length distributions for this isotype, which were similar to the data we observed in the absence of the KLC (Sup. Fig. 3, Fig. 2L-M). From these data, we conclude that the unique mechanochemical properties of KIF5A we observe in tail-truncated (Fig. 1) or full-length KHC complexes (Fig. 2), also extend to tetrameric kinesin motor complexes. However, we cannot rule out that the observed motile fraction of KIF5A motors was composed of KHC molecules that dissociated from the KLC. These data reveal that KLC-mediated motor inhibition strongly synergizes with the tail-mediated inhibition for all human KHC isotypes.

### KLC-Mediated Autoinhibition is Independent of Tail-Mediated Autoinihibiton

Previous studies on tail-mediated KHC autoinhibition utilized different types of mutations that are thought to bypass the molecular mechanism responsible for motor inhibition by the tail domain (Coy et al., 1999; Friedman and Vale, 1999; Kaan et al., 2011; Kelliher et al., 2018). First, deletion of a predicted break in the coiled-coil region of the stalk, termed the ‘hinge’ was shown to result in higher microtubule-stimulated ATPase activity (Coy et al., 1999; Friedman and Vale, 1999) and a variable increase in the number of processive molecules observed in a single-molecule assay (Friedman and Vale, 1999). The interpretation of these results was that without the hinge region, the KHC could not fold back on itself to allow an interaction between the C-terminal IAK domain and the N-terminal motor domains. However, evidence of kinesin unfolding was lacking in these studies. Contradictory data was obtained showing an apparent increase in sedimentation value for delta-hinge kinesin constructs, indicating potential oligomerization of the motor driven by the deletion (Coy et al., 1999). An atomic structure of dimeric *Drosophila* kinesin motor domains bound to a peptide that encompasses the IAK motif revealed important conserved residues within the motor domain that participate in the interaction with the tail domain (Fig. 2A, (Kaan et al., 2011)). Disruption of these interactions by mutation of these critical residues has been reported to disrupt kinesin autoinhibition in cells (Cai et al., 2007), enzymatic assays (Hackney and Stock, 2000), and in single molecule measurements performed in cell lysates (Kelliher et al., 2018). Additionally, the KLC has been reported to contribute to kinesin autoinhibition in a manner that is distinct from the IAK-mediated kinesin repression in cells (Cai et al., 2007).

We generated some of these previously reported mutations in KIF5B (Fig. 4A-B) in an effort to examine their effects on motor activity under identical assay conditions. We also purified recombinant KLC1, and sub-fragments encompassing the N-terminal α-helical region and C-terminal TPR domains (Fig. 4A-B), in order to directly observe how the KLC affects each type of autoinhibition mutant. We assessed the behavior of the mutant kinesin dimers in our single-molecule TIRF assay (Fig. 4C), and surprisingly observed very little processive movement of the mutant dimers (Fig. 4C-D). These observations are in contrast to a previous report of an increased fraction of processive delta-hinge mutant motors in a single-molecule assay (Friedman and Vale, 1999), although the increase in processive motors was variable in that study. In addition, the delta-hinge mutant, as well as mutation of residue K944 within *Drosophila* kinesin IAK domain (equivalent to K922 in human KIF5B), reportedly increased the velocity and distance traveled between pauses of single kinesin molecules in cell lysates (Kelliher et al., 2018). These observations were interpreted as evidence for decreased efficiency of tail-mediated autoinhibition, although an increase in the fraction of processive motors was not reported. Strikingly, we did not observe an increase in the fraction of processive motors when we assessed two equivalent mutations (Δhinge and IAKàAAA) in human KIF5B (Fig. 4C-D). Finally, mutation of the equivalent residues of H136 and D185 in the *Drosophila* kinesin motor prevented tail-peptide mediated inhibition of the motor’s enzymatic activity (Kaan et al., 2011), again interpreted as disrupting the tail-mediated inhibition of the motor domains. In contrast to these prior observations, we did not detect an increase in the fraction of processive mutant motors, as compared to wild-type kinesin dimers (Fig. 2K) in our purified system using the equivalent mutations H129A and E178A in human KIF5B (Fig. 4C-D). Rather, we instead observed a consistently large, ~50-100-fold increase in the microtubule landing rate of all of the mutant motors assayed (Fig. 4E), in agreement with previous notions that the IAK domain inhibits the microtubule association of the motor (Cai et al., 2007; Stock et al., 1999). From these data, we conclude that the main effect of abolishing the IAK-motor interaction is a dramatic increase in the microtubule-association rate. Our data therefore suggest that other elements within the mutant motors keep the motor in a conformation incompatible with processive movement.

**Figure 4.**
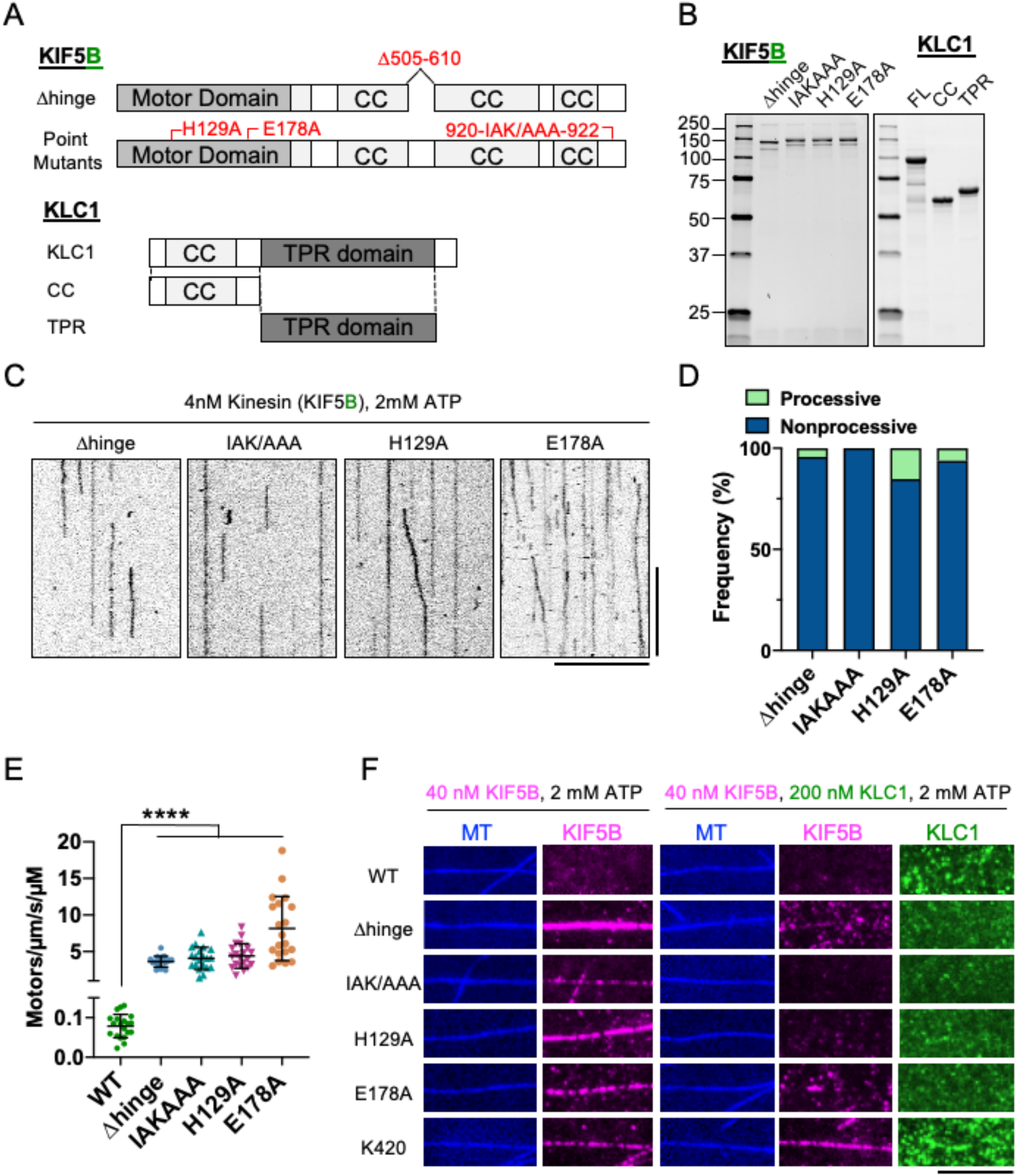
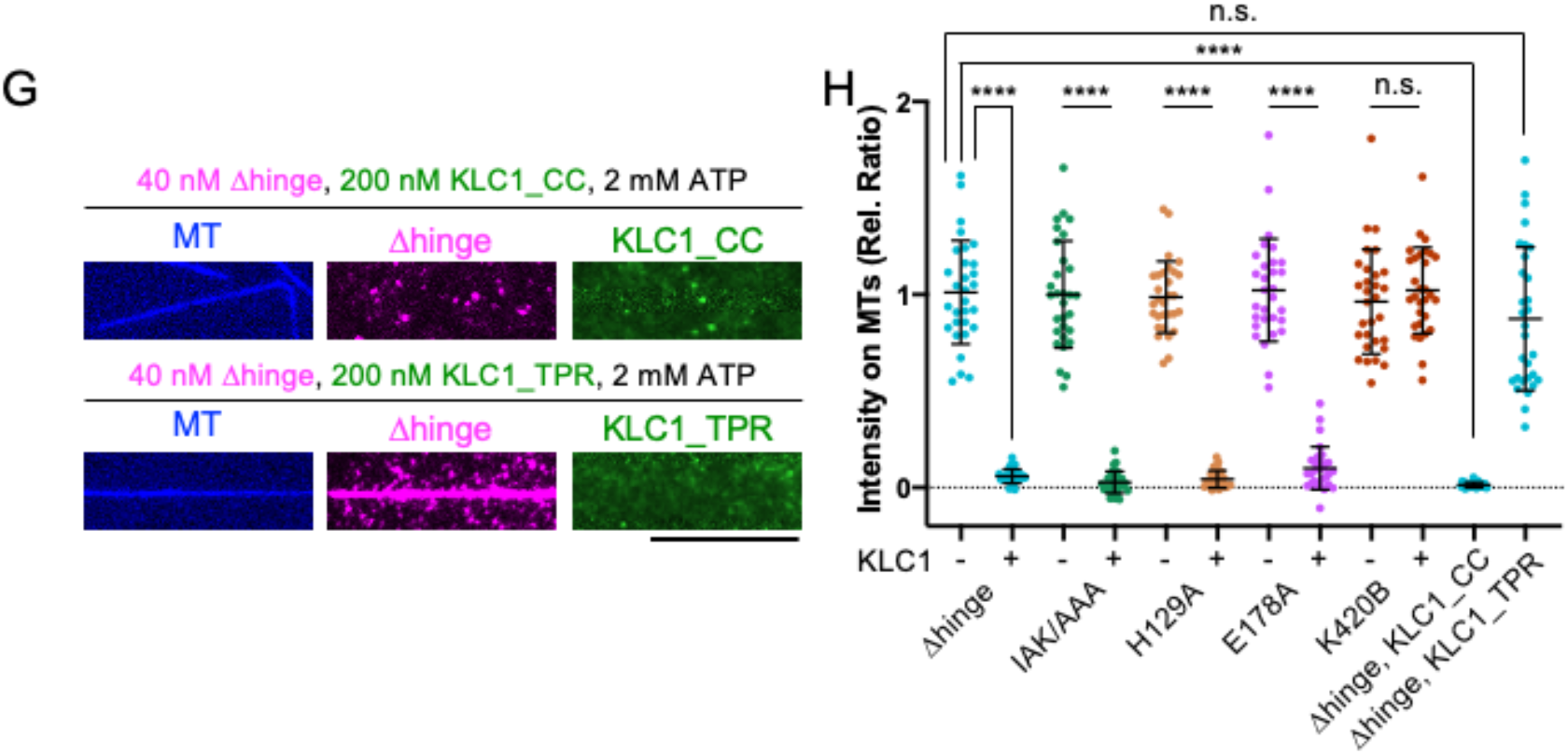
The KLC inhibits KIF5 independently of of the tail-inhibition mechanism. Schematic of KIF5B-Δhinge, IAK/AAA, H129A, E178A mutants and KLC1 and fragments used. KIF5B-Δhinge lacks aa 505-610. KLC1-CC or KLC1-TPR comprises aa 1-200 or aa 201-500 of human KLC1. (B) Coomassie blue stained gel showing the purified mScarlet-tagged KIF5 mutants and sfGFP-tagged recombinant KLC1. (C) Kymographs of KIF5 mutants on microtubules in the presence of 2 mM ATP. Scale bars are 10 s (vertical) and 10 μm (horizontal). (D) Measured frequency of processive and non-processive (static or diffusive) events. Processive runs were 4% for KIF5B-Δhinge, 0% for KIF5B-IAKAAA, 15% for KIF5B-H129A and 6% for KIF5B-E178A. N =2, n = 20 MTs, n = 280, 325, 367 and 645 molecules, respectively. (E) Measured landing rates of KIF5 mutants. Lines: mean ± SD: 0.08 ± 0.03 /μm/s/μM (for KIF5B-WT, the data of Fig. 2J is replotted here), 3.6 ± 0.7 /μm/s/μM (KIF5B-Δhinge), 4.1 ± 1.5 /μm/s/μM (KIF5B-IAKAAA), 4.4 ± 1.7 /μm/s/μM (KIF5B-H129A), 8.1 ± 4.4 /μm/s/μM (KIF5B-E178A). N = 2, n = 20 MTs, respectively. One-way ANOVA followed by Tukey’s multiple comparison test was performed. **** Adjusted P < 0.0001; n.s., not significant. (F) TIRF images of 40 nM KIF5B (magenta) on microtubules (blue) with or without 200 nM recombinant full-length KLC1 (green) in the presence of 2 mM ATP. Scale bar is 10 μm. (G) TIRF images of 40 nM KIF5B-Δhinge (magenta) on microtubules (blue) with 200 nM recombinant KLC1 (KLC1-CC or KLC1-TPR) in the presence of 2 mM ATP. Scale bar is 10 μm. (H) Quantification of relative fluorescence intensity of KIF5B on MTs in Fig. 4F or 4G. Lines: mean ± SD. In each mutant, fluorescence intensities were normalized to the average intensity of KIF5B in the absence of KLC1. N = 30. For KIF5B-IAKAAA, KIF5B-H129A, KIF5B-E178A and K420B, two-tailed Student’s t-tests were performed. **** P < 0.0001; n.s., not significant. For KIF5B-Δhinge, One-way ANOVA followed by Tukey’s multiple comparison test was performed. **** Adjusted P < 0.0001; n.s., not significant.

The KLC, or its N-terminal subdomain, has been reported to strongly inhibit the microtubule association of the KHC in cell lysates (Verhey et al., 1998), and in purified systems (Coy et al., 1999; Friedman and Vale, 1999). FRET measurements in cells suggested that the C-terminal TPR domain of the KLC (Fig. 4A) was necessary to push the kinesin motor domains apart into a conformation that was incompatible with motility (Cai et al., 2007). In order for this to occur, the tail region of the motor must be able to fold, presumably at the hinge region, to allow proximity of the KLC TPR domains to the motor domains. To test this idea, we added purified KLC domains to KHC motors in trans and assessed the ability of the KHC to bind to microtubules in the presence of ATP. As we previously observed (Fig. 2), wild-type KHC had a low apparent affinity for microtubules (Fig. 4F, H), while in agreement with our landing rate measurements (Fig. 4E), the mutant KHC motors all showed substantially higher microtubule association (Fig. 4F, H) in the absence of the KLC. When a five-fold molar excess of purified KLC was included, we observed strong attenuation of the microtubule association of all of the KHC motors. This result is very surprising given that all the current models for kinesin autoinhibition propose that the tail must be able to fold back to allow the IAK domain to bind to the motors, and also to place the KLCs in close proximity to the motor domains (Cai et al., 2007; Coy et al., 1999; Dietrich et al., 2008; Friedman and Vale, 1999). In theory, this should not be possible when the hinge region is removed, or when critical interaction residues between the tail and motor domains are abolished. Purified KLC did not directly bind to microtubules, and it did not obviously affect the microtubule association of tail-truncated KIF5B (Fig. 4F, H), suggesting that the KLC does not block motors from binding microtubules non-specifically in our assay. Finally, we assessed which subdomain of the KLC was necessary for the strong inhibitory effect we observed. In agreement with prior findings in cell lysates (Verhey et al., 1998), we found that the N-terminal coiled-coil containing region, but not the isolated TPR domains, was sufficient to strongly attenuate the microtubule association of the delta-hinge KHC mutant (Fig. 4G-H). Thus, the small N-terminal region of the KLC is sufficient to strongly repress microtubule binding of the KHC even in the absence of a direct interaction between the tail and motor domains. Prior models for the mechanism of kinesin autoinhibition are insufficient to account for these observations.

### Binding of a W-acidic Motif Cargo-Adapter to the KLC is Sufficient to Relieve Kinesin Tetramer Autoinhibition

Previous work delineated two classes of KLC binding peptides found in a range of molecules termed W-acidic, and Y-acidic, due to highly conserved tryptophan- or tyrosine-containing motifs that directly interact with the TPR domains of the KLC (Cross and Dodding, 2019). Attachment of the W-acidic peptide to intracellular cargo results in kinesin-dependent displacement of cargo in cells (Kawano et al., 2012). However, it remains undetermined if binding of the W-acidic peptide to the KLC within the kinesin tetramer is sufficient to activate the autoinhibited tetramer, or if other cellular factors participate in motor activation. To examine if the binding of W-acidic motifs activates the motor, we first generated a recombinant kinesin adapter protein from the nesprin family of trans-nuclear envelope proteins that are known to contain a conserved ‘LEWD’ motif that binds directly to the KLC (Roux et al., 2009; Wilson and Holzbaur, 2015). Because many members of the nesprin family are very large, we chose one of the smallest family members that contains the ‘LEWD’ motif that was previously shown to interact with kinesin: nesprin-4 (Roux et al., 2009). We purified the cytoplasmic domain of nesprin-4 (Fig. 5A), and confirmed that it interacted with endogenous kinesin from rat brain lysates in a pulldown assay (Fig. 5B). Importantly, mutation of the critical ‘LEWD’ sequence to ‘LEAA’ completely abolished kinesin binding, as expected (Fig. 5B, (Roux et al., 2009; Wilson and Holzbaur, 2015)). Thus, purified nesprin-4 can interact with endogenous kinesin complexes.

**Figure 5.**
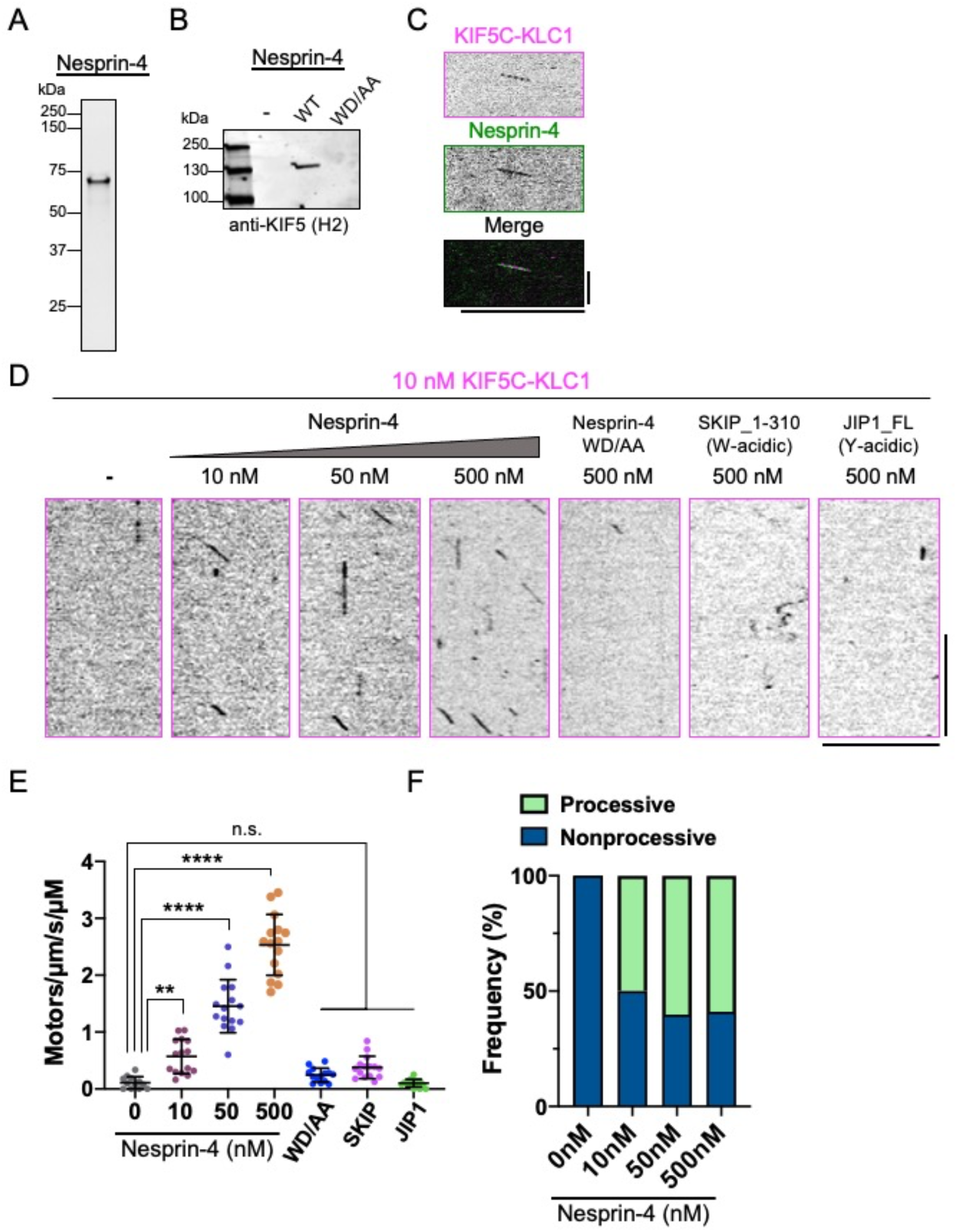
Nesprin-4 activates the autoinhibited KIF5-KLC tetramer. **(A)** Coomassie blue stained gel showing purified sfGFP-tagged cytosolic domain of human Nesprin-4. **(B)** Pull-down assay with Nesprin-4 (wild-type and WD/AA) from rat brain lysate. **(C)** Kymograph from multi-color TIRF movies showing co-movement of of mScarlet-tagged KIF5C-KLC1 (magenta) tetramer with sfGFP-tagged nesprin-4 (green) on microtubules in the presence of 2 mM ATP. 10 nM of each protein was used. Individual fluorescent channels (top and middle) and the merged kymograph (bottom) are shown. Scale bars are 10 s (vertical) and 10 μm (horizontal). **(D)** Kymographs showing the behavior of mScarlet tagged KIF5C-KLC1 mixed with indicated proteins in the presence of 2 mM ATP. Scale bars are 10 s (vertical) and 10 μm (horizontal). **(E)** Measured landing rates of KIF5-KLC1 tetramers. Lines show mean ± SD: 0.1 ± 0.1 /μm/s/μM (Note, for 10 nM KIF5C-KLC1 alone, the data from Fig. 3I was plotted again), 0.6 ± 0.3 /μm/s/μM (10 nM KIF5C-KLC1 + 10nM nesprin-4), 1.5 ± 0.5 /μm/s/μM (10 nM KIF5C-KLC1 + 50 nM Nesprin-4), 2.5 ± 0.5 /μm/s/μM (10 nM KIF5C-KLC1 + 500 nM Nesprin-4), 0.2 ± 0.1 /μm/s/μM (10 nM KIF5C-KLC1 + 500 nM Nesprin-4_WDAA), 0.4 ± 0.2 /μm/s/μM (10 nM KIF5C-KLC1 + 500 nM SKIP_1-310), 0.1 ± 0.1 /μm/s/μM (10 nM KIF5C-KLC1 + 500 nM JIP1). N = 2. n = 12, 15, 15, 15, 14, 15 and 15 microtubules respectively. One-way ANOVA followed by Dunnett’s multiple comparison test was performed. ** Adjusted P < 0.01; **** Adjusted P < 0.0001; n.s., not significant. **(F)** Frequency of processive and non-processive (static or diffusive) events. Processive runs were 0%, 24% 50% and 60% for 0, 10, 50 and 500 nM Nesprin-4. n = 89, 17, 112 and 221 molecules. The data of Fig. 3J was plotted again for 0 nM nesprin-4.

Next, we combined recombinant KIF5C-KLC1 tetramer with purified nesprin-4 in our single molecule assay. W-acidic peptides have low micromolar affinities for isolated KLC TPR domains (Pernigo et al., 2013), and the native KLC is reportedly autoinhibited (Yip et al., 2016), making observation of single molecule interactions at low nanomolar concentrations difficult. Nonetheless, at 10 nM concentration of each protein, we observed infrequent processive events that clearly contained both KHC-mScarlet and sfGFP-nesprin-4 signals, confirming a direct interaction between the kinesin tetramer and nesprin-4 (Fig. 5C). We observed that 77% of processive kinesin molecules contained nesprin-4 signal, further revealing that most motile kinesin complexes were bound to nesprin-4. The addition of higher concentrations of nesprin-4 obscured the nesprin-4 signal due to high background, but we observed a clear dose-dependent increase in the frequency of kinesin landing (0.11 ± 0.1, 0.6 ± 0.3, 1.5 ± 0.5, 2.5 ± 0.5 motors/μm/s/μM (s.d) for 0, 10, 50, and 500 nM nesprin-4, respectively. Fig. 5D-E). Of these landing events, we observed a significant increase in the percentage of processive kinesin events (Fig. 5F). In agreement with our pulldown data (Fig. 5B), we did not observe any increase in landing events in the presence of a large excess of the nesprin-4 mutant containing a mutated W-acidic motif (Fig. 5D-F). The velocity and run-lengths of the activated kinesin tetramers were similar across nesprin-4 concentrations (Fig. S2), suggesting the cargo-activated tetramer has similar biophysical properties to the uninhibited motors (Fig. 1). Thus, in a purified system, the binding of a W-acidic motif adapter molecule to the kinesin tetramer is sufficient to relieve motor autoinhibition.

We tested an N-terminal fragment of the kinesin adapter protein SKIP, which also contains W-acidic motifs and binds to the isolated KLC TPR domain in vitro (Pernigo et al., 2013). Despite the well characterized, albeit weak, binding of SKIP to the isolated KLC TPR domain, we did not observe any effect of this fragment in our single molecule assay (Fig. 5D-E). We additionally tested an orthogonal adapter protein, JIP1, which contains a Y-acidic motif, and has been shown to bind directly to the KLC TPR domain (Pernigo et al., 2018), and suggested to activate KHC motility in cell lysates (Fu and Holzbaur, 2013). We again did not observe any effect of JIP1 on the frequency of processive events with our purified kinesin tetramer (Fig. 5D-E). These results reveal that distinct adapters have differential abilities to activate the motor in vitro. We speculate that binding affinity, and potentially unexplored autoinhibition of the adapter molecules, likely play key roles in the activation process. Further work is needed to carefully dissect the interactions of distinct adapter proteins with the intact tetrameric kinesin motor complex.

### MAP7 Enhances Kinesin Tetramer Activation

Despite the effect of nesprin-4 on the autoinhibited kinesin tetramer, we noticed that the landing rate of the activated kinesin-nesprin complexes (Fig. 5E) was nonetheless ~ 50-fold lower than that of the tail-truncated dimers (Fig. 1D). This observation suggests that, while the inhibition of motor activity is relieved by the binding of an activating adapter, the full potential of the kinesin-microtubule interaction remains stifled. Interestingly, recent work has uncovered a role for the non-motor microtubule-associated protein 7 (MAP7) family of proteins in kinesin motility. In vitro, MAP7 proteins dramatically enhance the binding of kinesin to the microtubule lattice, and also potentially affect the autoinhibition state of the dimeric motor (Hooikaas et al., 2019; Monroy et al., 2018). In living cells, kinesin activity is strongly recruited to microtubules enriched with MAP7 proteins (Serra-Marques et al., 2020), suggesting that recruitment of activated kinesin motors to microtubules by MAP7 proteins plays a central role in kinesin motility in organisms that contain MAP7 homologues.

We reasoned that the relatively low microtubule-association rate of activated kinesin motors could be due to the lack of a MAP7 family protein in our reconstitution. A prior study found that the kinesin-binding domain of MAP7 could increase the landing rate of tail-truncated kinesin (amino acids 1-560), suggesting that elements outside of the kinesin tail domain may also contribute to motor regulation (Hooikaas et al., 2019). However, any direct consequence of MAP7 on the autoinhibition mechanism of fulllength kinesin tetramer remains unexplored. First, we examined the effects of MAP7 on the autoinhibited KIF5C-KLC1 tetramer and observed a strong increase in the number of kinesin tetramers binding to microtubules (Fig. 6A, compare to Fig. 3H). Although the slower frame rates at which we acquired these data (see below) preclude a direct quantitative comparison with our data of the kinesin tetramer alone (Fig. 3), the presence of MAP7 on the microtubule clearly increased the overall amount of the KIF5C-KLC1 tetramer on the microtubule. However, MAP7 did not obviously lead to the activation of processive motility, as we very rarely observed processive movement of the bound KIF5C-KLC1 tetramers (Fig. 6A). Thus, we conclude that MAP7 strongly recruits the full kinesin tetramer to microtubules, but does not relieve the autoinhibition of the tetrameric complex.

**Figure 6.**
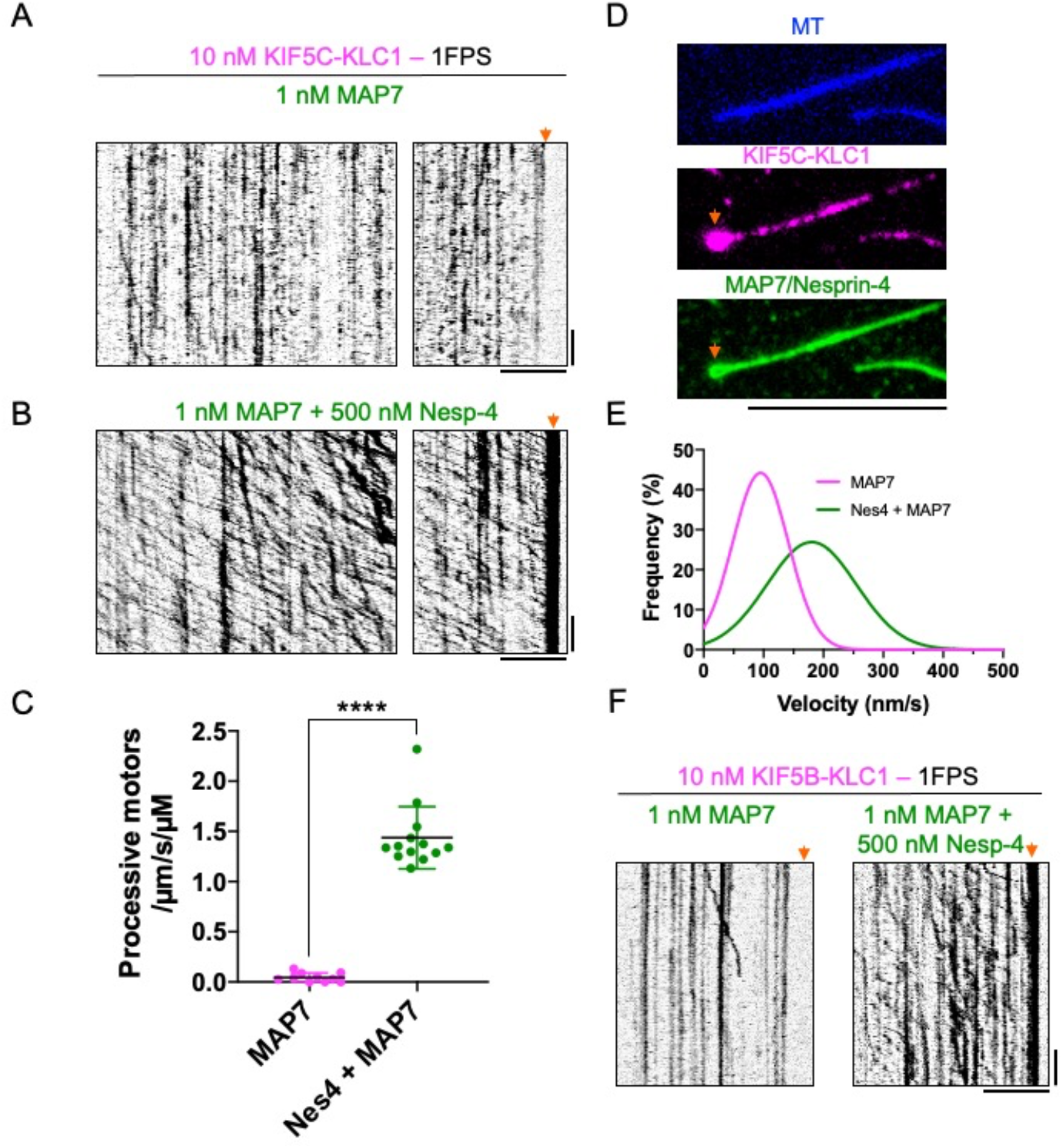
Synergistic activation of kinesin tetramers by nesprin-4 and MAP7. **(A)** Kymographs of KIF5C-KLC1 on MAP7-coated microtubules. Note the lack of processive movement. Arrow highlights no accumulation of motors at the microtubules ends. **(B)** Kymographs of KIF5C-KLC1 on MAP7-coated microtubules in the presence of nesprin-4. Note the highly processive movement. Arrow highlights strong accumulation of processive motors at one end of the microtubule. Scale bars are 30 s (vertical) and 5 μm (horizontal). **(C)** Measured landing rates of processive KIF5C-KLC1 motors in the absence or presence of MAP7. Only processive molecules were counted in this analysis. Lines: mean ± SD: 0.045 ± 0.045 /μm/s/μM (KIF5C-KLC1 with MAP7), 1.4 ± 0.31/μm/s/μM (KIF5C-KLC1 with MAP7 and Nesprin-4). N = 2. n = 10 or 13 MTs, respectively. Two-tailed Student’s t-tests were performed. **** P < 0.0001. **(D)** TIRF images of KIF5C-KLC1 (magenta, middle) accumulation at one microtubule end (arrows) in the presence of MAP7 and nesprin-4 (both green, bottom). Note green accumulation at the microtubule end is presumably nesprin-4 signal transported there by kinesin motors. Scale bar is 10 μm. **(E)** Gaussian fits of KIF5C-KLC1 velocities distributions in the presence of MAP7 or MAP7 + nesprin-4. Mean ± SD: 95 ± 47 nm/s (MAP7, magenta), 180 ± 75 nm/s (MAP7 + Nesprin-4, green). N = 2. n = 21 or 706 molecules, respectively. **(F)** Kymographs of KIF5B-KLC1 on MAP7-coated microtubules with (right) or without (left) nesprin-4. Arrows indicate MT ends. Note accumulation of processive motors at microtubule ends only in the presence of MAP7 and nesprin-4. Scale bars are 30 s (vertical) and 5 μm (horizontal).

Next, we added both nesprin-4 and MAP7 to the assay to assess the compound effects of nesprin-4-mediated release of autoinhibition and MAP7 recruitment of tetrameric motors. Inclusion of both molecules in the assay resulted in a striking enhancement of robustly processive movement of kinesin tetramers along microtubules (Fig. 6B). The presence of MAP7 stimulated the landing of processive kinesin motors by over 30-fold (Fig. 6C). The motility of nesprin-4-activated KIF5C-KLC1 tetramers in the presence of MAP7 was dramatically distinct from the behavior of nesprin-4 activated motors alone (compare Figs. 6B and 5D). In the presence of MAP7, activated motor movement was highly processive, with run-lengths that clearly exceed 5 μm (Fig. 6B). Such long runs were never observed in the absence of MAP7. Indeed, we noted a strong accumulation of motors at the presumed plus-end of the microtubule (Fig. 6B, C), suggesting that many motors moved until they reached the end of the filament. The apparently weak affinity between nesprin-4 and kinesin precluded the dilution of motors in the assay, and the resulting high density of motors recruited to the microtubule by MAP7 made accurate assignment of the beginning and endings of runs difficult. We therefore measured uninterrupted segments of processive movements in the kymographs and found that in the presence of MAP7, the characteristic run-length of KIF5C-KLC1 was at least 3.5 μm, approximately 7-fold longer than we observed for nesprin-activated tetramers in the absence of MAP7 (Fig. S2). We note that this value is an underestimate of the true run-length due to the limitations of motor concentration in the assay. Differences in image acquisition rates precluded a direct quantitative comparison with the measured run-lengths of nesprin-4 activated motors in the absence of MAP7 (Fig. S2), but the stimulation of kinesin tetramer processivity we observe in the presence of MAP7 is in agreement with previously reported, though more modest, increases in the processivity of tail-truncated or full-length dimeric kinesin (Hooikaas et al., 2019; Monroy et al., 2018). We also measured the velocity of the rare processive molecules in the presence of MAP7 and the processive molecules in the presence of both MAP7 and nesprin-4. The velocity of processive kinesin in the presence of MAP7 was low (95 ± 46.5 nm/sec and 180 ± 74 nm/sec (s.d.) for MAP7 and MAP7 with nesprin-4 respectively, Fig. 7E), also in agreement with prior work showing MAP7 hinders kinesin velocity (Ferro et al., 2020; Hooikaas et al., 2019; Monroy et al., 2018). Finally, the synergy of MAP7 and nesprin-4 in our assay was also observed with the ubiquitously-expressed KIF5B-KLC1 isotype of kinesin (Fig. 6F), revealing a general pathway for kinesin activation through the binding of a cargo-adapter molecule to the light chains and recruitment of the activated complex to microtubules via MAP7.

## Discussion

The kinesin-1 motor is the most ubiquitous and well-studied plus-end directed transport kinesin in biology. Despite decades of research on the mechanism of kinesin mechanochemistry, we still lack a fundamental understanding about how cells regulate kinesin motor activity for the correct spatiotemporal delivery of plus-end directed cargos. Additionally, humans express a diverse set of KHC and KLC isotypes, presumably resulting in the ability to fine tune motor activity for specific cellular functions. If and how these distinct kinesin isotypes differ from each other remains unclear. In this study, we have systematically studied the different KHC isotypes expressed in humans. We first delineated isotype-specific differences in the mechanochemistry of truncated, uninhibited motors. We find that despite high sequence homology, KIF5A motors are uniquely fast among the three human isotypes, yet further work is required to precisely determine the kinetic steps that vary between motor isotypes. Our results suggest that in the absence of the KLC, the KIF5A heavy chain has a tendency to oligomerize into tetramers (Figs. 2-3). This result may have implications for human disease, as mutations in the divergent KIF5A tail domain cause ALS (Nicolas et al., 2018).

Tail-mediated autoinhibition is the most widely studied mechanism of kinesin regulation (Cross and Dodding, 2019; Dietrich et al., 2008; Hackney and Stock, 2000; Stock et al., 1999). We find evidence that tail-mediated repression of motor activity varies by KHC isotype. Despite conservation of the IAK motif in all KIF5 genes, tail-mediated repression of motor activity is weakest for KIF5A, which may be related to its divergent C-terminal extension. The tendency of KIF5A to oligomerize in the absence of the KLC could also contribute to its uniquely high intrinsic activity (Fig. 2). It is currently unclear what fraction of the total cellular pool of KHC operates in the absence of the KLC. Certain functions of kinesin motors may be independent of the KLC (Palacios and St Johnston, 2002), such as the transport of mitochondria (Glater et al., 2006; van Spronsen et al., 2013), although the data for this assertion are conflicting (Khodjakov et al., 1998). Thus, a more thorough understanding of the effects of the divergent kinesin tail domains on the autoinhibition mechanism of the KHC dimer will inform on cargo-specific regulation of kinesin motor activity.

Consistent with prior work (Verhey et al., 1998), we find that association of the KLC with the KHC leads to a fully repressed motor that is incapable of productive interactions with the microtubule. While Verhey et al. (Verhey et al., 1998) suggested the distal tail region of KHC is required for the KLC-mediated regulation, our data show that the IAK domain within this segment of the KHC tail is not involved in KLC-mediated regulation. Binding of the KLC does not prevent motor association with microtubules in the presence of an ATP analogue, implying an allosteric mechanism by which the KLC blocks one or more of the required steps in the kinesin mechanochemical cycle, as opposed to physically blocking the motor from binding the microtubule. Further work is necessary to precisely define the step(s) in the kinesin mechanochemical cycle that are altered by the KLC. Interestingly, only a small piece of the KLC N-terminus is required for this large effect ((Verhey et al., 1998), Fig. 4). Previous data have suggested a model whereby the kinesin molecule folds in half to allow direct interactions between the motor and tail domains (Cai et al., 2007; Coy et al., 1999; Friedman and Vale, 1999; Kaan et al., 2011; Stock et al., 1999). A folded conformation would place the KLC adjacent to the motor domain, possibly facilitating direct effects. However, we found that removing the previously described hinge region, or mutating residues important for this putative motor-tail interaction, did not prevent the KLC from exerting its inhibitory effect on the motor-microtubule interaction. The current intramolecular folding model cannot account for these data. Since the binding site for the KLC is separated from the motor domain by ~ 450 amino acids, we hypothesize that a long-range allosteric mechanism could account for such an effect. This proposed mechanism could propagate conformational changes down the central stalk region of kinesin upon KLC association.

Our data with KHC dimers is also not fully consistent with the current model for kinesin autoinhibition. Several prior studies have suggested that removal of either the central hinge region or the IAK domain lead to motor activation (Coy et al., 1999; Friedman and Vale, 1999; Kelliher et al., 2018). However, some of these studies did not directly assay motor movement as we have done here. Using single molecule analysis, Friedman and Vale reported an increase in the frequency of processive KHC dimers when the hinge region was deleted (Friedman and Vale, 1999). However, this effect varied from 5-50-fold across the two protein preparations reported. In addition, the velocity of these motors (~300 nm/sec) was three-fold slower than that of the motors activated by nesprin-4 reported here, possibly due to differing assay conditions. Kelliher et al. (Kelliher et al., 2018) reported that deletion of the hinge domain, or mutation of analogous residues important for the interaction between the motor domain and IAK motif in *Drosophila* kinesin-1 led to an increase in motor velocity and run-length between run pauses. These results were interpreted as evidence of disruption of KHC autoinhibition; however, the study did not compare the landing frequency of motile motors. Our results with similar mutations in the human KHC suggest the primary effect may be to increase the landing frequency of motors onto the microtubule. If a fraction of motors is inherently active, increasing the landing rate could result in more observed motile motors in total without substantially changing the proportion of active motors. An increased landing rate onto microtubules could also account for the observed enhancement of kinesin enzymatic activity upon disruption of the central hinge region, or motor-IAK interactions (Coy et al., 1999; Kaan et al., 2011), if the motors turn over ATP without directly coupling it to processive stepping. Consistent with this hypothesis, we observe that mutant KHC motors bind and release from the microtubule in the presence ATP (Fig. 4). Thus, our single molecule data revise the current models for KHC autoinhibition, but a coherent molecular mechanism remains obscured by the lack of available structural data. High-resolution structures of the autoinhibited kinesin dimer and tetramer would greatly inform on these questions, and should be a high priority for future studies.

Using purified components, we have demonstrated that binding of a cellular cargo adapter, nesprin-4, to the kinesin tetramer is sufficient to relieve one facet of the autoinhibition mechanism of the motor. Nesprins are nuclear envelope proteins that are well-established to play roles in linking the nucleus to the major cytoskeletal networks through direct interactions (Fridolfsson et al., 2018), and interactions with cytoskeletal motors. Different nesprin family members contain consensus W-acidic motifs that directly bind to the TPR domains of the KLC (Pernigo et al., 2013). Our data reveal that the binding of the W-acidic motif to the intact KLC within the kinesin tetramer is sufficient to relieve the KLC-induced inhibition of motor activity. However, this interaction is weak and inefficient in vitro, and further studies are necessary to determine why. One likely reason is the previously reported autoinhibition of the KLC itself (Yip et al., 2016). However, other well-characterized cargo adapters such as SKIP contain similar KLC-binding motifs, but a SKIP fragment containing these motifs did not suffice to activate the motor in our reconstitution for reasons that remain unclear (Fig. 5). One clue may be related to the architecture and native cellular environment of the kinesin adapter. Nesprins are thought to act as trimers within the outer nuclear envelope (Sosa et al., 2013) and this architecture raises questions about the stoichiometry of interaction with the kinesin dimer, as each nesprin-4 molecule contains only one W-acidic motif. On the other hand, SKIP contains tandem W-acidic motifs, and associates with membranes peripherally (Pernigo et al., 2013). The oligomeric state of SKIP is currently unknown, and we suggest that the oligomerization state, and potentially the native environment within or near a membrane could play key roles in the effectiveness of motor activation by kinesin cargo-adapter molecules. Similarly, the Y-acidic motif-containing molecule, JIP1, was insufficient to activate the motor in our assay, consistent with prior results in cell lysates (Blasius et al., 2007; Kawano et al., 2012). This work also reported that JIP1 coordinates with another molecule, FEZ1, that binds directly to the HC, to activate the kinesin tetramer in cell lysates. These results again suggest that each cargo adapter will require specific characterization to uncover novel regulatory schemes that may control its ability to activate kinesin tetramer motility. The molecular mechanisms described here are reminiscent of the cargo-adapter specific effects of dynein-dynactin activation (Reck-Peterson et al., 2018) and further work is necessary to uncover if autoinhibition, oligomerization, and post-translational modifications play key roles in controlling the activity of kinesin cargo-adapters. The in vitro reconstitution assay we have developed here will no doubt play a key role in these future efforts.

It is striking that the activated kinesin-nesprin complex is still approximately 75-150-fold less efficient at initiating movement along microtubules as compared to truncated, constitutively active kinesin molecules (Fig 1D vs 5E). This result suggests that cargo-adapter binding may not lead to full activation of the motor, which is consistent with the existence of distinct KLC- and KHC-dependent mechanisms of autoregulation. Cargo-binding to the KLC can unlock one of them but additional factors may be required for full activation. The non-motor MAP7 protein strongly stimulates kinesin landing onto microtubules (Ferro et al., 2020; Hooikaas et al., 2019; Monroy et al., 2018; Sung et al., 2008). MAP7 proteins have also been suggested to activate the kinesin dimer (Hooikaas et al., 2019), but a direct measurement of the fraction of active motors was not presented in this study. Intriguingly, binding of a MAP7 fragment to the stalk domain of a truncated kinesin could still enhance kinesin’s landing rate, even in the absence of a direct MAP7-microtubule interaction (Hooikaas et al., 2019). This result implies that the stalk domain of kinesin exerts regulatory control over the kinesin-microtubule interaction, and that proteins such as MAP7, that directly interact with this region may exert allosteric control over kinesin motor activity. This model of kinesin regulation is distinct from the folding model currently prevalent in the field, and is consistent with our data using kinesin mutants to prevent the head-tail interaction. We propose a model whereby long range allostery could propagate conformational changes down the central stalk region to control kinesin motor activity. In line with prior results, we found that the inclusion of MAP7 into our reconstitution greatly enhanced the landing of activated kinesin molecules onto microtubules and strongly enhanced their run-lengths (Ferro et al., 2020; Hooikaas et al., 2019; Monroy et al., 2018). MAP7 alone, however, did not lead to robust activation of the kinesin tetramer, implying a synergistic relationship requiring cargo-adapter binding to the light chains to relieve inhibition of motor activity, and a direct interaction with a non-motor MAP to facilitate recruitment to microtubules. We note there is also strong genetic evidence for synergy between kinesin and MAP7 *in vivo* (Metzger et al., 2012). Not all organisms that contain a kinesin-1 homologue also contain obvious MAP7 homologues. It will be interesting to examine the activation mechanisms of these kinesin-1 homologues in the future.

We are struck by the parallels of this model to the recently worked out cytoplasmic dynein regulatory scheme, whereby a cargo-adapter molecule is required to facilitate the formation of a complex between dynein and its activator dynactin (McKenney et al., 2014; Schlager et al., 2014). Dynactin itself acts as a non-motor MAP, that directly facilitates the landing of the activated dynein-dynactin-cargo-adapter complex onto specific subsets of tyrosinated microtubules (McKenney et al., 2014; McKenney et al., 2016). In cells, MAP7 appears to target kinesin activity to subsets, or subregions, of microtubules (Serra-Marques et al., 2020), and future work focused on determining the mechanisms that target MAP7 to microtubules will no doubt shed light on how cells regulate the spatiotemporal activity of the kinesin motor.

In summary, we have reported intrinsic biophysical differences between human KHC isotypes and reconstituted a two-factor activation mechanism of the human kinesin-1 tetramer. We have found that binding of a cargo-adapter protein to the KLC is sufficient to relieve an autoinhibitory mechanism that blocks motor activity; however, the removal of this blockade does not induce robust motor motility along microtubules. We found that the activated kinesin tetramer must further be recruited to microtubules by the non-motor MAP7, which greatly enhances both the landing rate and processivity of the tetrameric complex. Our results have broad implications for understanding how cells control intracellular transport, a process that underlies a growing list of human diseases.

## Materials and Methods

### Plasmids

A cDNA plasmid encoding human *KIF5A* (BC146670), *KIF5B* (BC126281), *KIF5C* (BC110287), *KLC1* (BC008881), *SKIP* (BC040441) were purchased from Transomics (Huntsville, AL, USA). Mouse *JIP1* cDNA (NM011162) was a gift from Dr. Toshiharu Suzuki (Hokkaido University, Japan). A human MAP7 plasmid was used as previously described (Monroy et al., 2018). A cDNA plasmid encoding human Nesprin-4 (NM_001297735, natural variant harboring Q165H) was a kind gift from Dan Starr (UC Davis). For preparing K420B, a DNA fragment encoding human KIF5B aa 1-420 was codon-optimized for E. coli and synthesized by gBlocks (Integrated DNA Technologies, Coralville, IA, USA).

Gibson assembly was used in the following constructs except as otherwise noted. For K420 series, KIF5A (aa 1-416), KIF5B codon optimized for E.coli (encoding aa 1-420 of human KIF5B as noted above), and KIF5C (aa 1-416) were cloned into pET28a vector with a C-terminal mScarlet-StrepII tag. The cytosolic domain of Unc-83c (1-698) was cloned into pET21a vector with C-terminal sfGFP-StrepII tag. Full-length KLC1, KLC1-CC (aa 1-200), KLC1-TPR (aa 201-500), cytosol domain of Nesprin-4 (aa 1-240), SKIP (1-310) and full-length JIP1 were cloned into pET28a vector with N-terminal 6xHis-2xStrepII-sfGFP-2xPrescission protease site. The WD/AA mutation was introduced into pET28a-Nesprin-4 using primers containing desired mutation which are complementary to opposite strands of the cDNA and internal primers of pET28a. For fulllength KIF5 dimers, full length KIF5s were cloned into pACEBac1 vector with a C-terminal mScarlet-StrepII tag. Using the pACEBac1-full-length KIF5B, KIF5B-ΔHinge (Δ505-610) was prepared using primers connecting C-terminus of KIF5B aa 1-504 with N-terminus of KIF5B aa 611-963 and internal primers of pACEBac1. KIF5B-IAK/AAA, H129A and E178A were prepared using primers harboring the desired mutations and internal primers of pACEBac1. For KIF5-KLC1 tetramer, KLC1 was first cloned into pIDS vector with N-terminal 6xHis-FLAG. pIDS-KLC1 vector was fused with either pACEBac1-KIF5A, KIF5B or KIF5C using Cre-Lox recombination. All constructs were verified by Sanger sequencing.

### Antibodies

Anti-KIF5 (H2, #MAB1614) was purchased from Sigma-Aldrich.

### Protein expression and purification

Bacterial expression and preparation of K420A, K420B, K420C, Nesprin-4, SKIP and JIP1 were performed as below. BL21-CodonPlus (DE3)-RIPL (Agilent) transformed with each plasmid were grown at 37°C in LB medium with Kanamycin until an optical density at 600 nm (OD_600_) reaches 0.4. The cultures were allowed to cool down to room temperature and induced by 0.2 mM isopropyl-β-D-thiogalactoside overnight at 18°C. Cells were harvested and resuspended in 25 ml of lysis buffer (50 mM HEPES-KOH, pH 7.5, 150 mM KCH_3_COO, 2 mM MgSO4, 1 mM EGTA, 10% glycerol) supplemented with 1 mM DTT, 1 mM PMSF, DNaseI and 0.1 mM ATP. Cell were lysted by passage through an Emulsiflex C3 high-pressure homogenizer (Avestin). Then the lysates were centrifuged at 15000 x g for 20 min at 4°C. The resulting supernatant were subject to affinity chromatography described below.

For constructs cloned into insect cell expression vectors (pACEBac1, pACEBac1/pIDS), DH10MultiBac (Geneva Biotech) were transformed to generate bacmid. Sf9 cells were maintained as a suspension culture in Sf-900II serum-free medium (SFM) (Thermo Fisher Scientific) at 27°C. To prepare baculovirus, 1 x 10^6^ cells of Sf9 cells were transferred to each well of a tissue-culture treated 6 well plate. After the cells attached to the bottom of the dishes, about ~5 μg of bacmid were transfected using 6 μl of CellfectinII reagent (Thermo Fisher Scientific). 5 days after initial transfection, the culture media were collected and spun at 3000 x g for 3 min to obtain the supernatant (P1). Next, 50 ml of Sf9 cells (2 x 10^6^ cells/ml) was infected with 50 μl of P1 and cultured for 5 days to obtain P2 viral supernatant. The resulting P2 were used for protein expression. For protein expression, 400 ml of Sf9 cells (2 x 10^6^ cells/ml) were infected with 4 ml of P2 virus and cultured for 65 hr at 27°C. Cells were harvested and resuspended in 25 ml of lysis buffer (50 mM HEPES-KOH, pH 7.5, 150 mM KCH_3_COO, 2 mM MgSO_4_, 1 mM EGTA, 10% glycerol) along with 1 mM DTT, 1 mM PMSF, 0.1 mM ATP and 0.5% TritonX-100. After incubating on ice for 10 min, the lysates were centrifuged at 15000 x g for 20 min at 4°C. The resulting supernatant were subject to affinity chromatography described below.

For affinity chromatography, the supernatants were pumped over a column of Streptactin XT resin (IBA) for ~1 hour at 4°C. The columns were then washed with excess lysis buffer to remove unbound material and the proteins were eluted in lysis buffer containing 100 mM D-biotin. Eluted proteins were further purified as described below.

K420A, K420B and K420C were further purified via anion exchange chromatography using a TSKgel SuperQ-5PW (Tosoh bioscience) 7.5 mm ID x 7.5 cm column equilibrated in HB buffer (35 mM PIPES-KOH pH 7.2, 1 mM MgSO4, 0.2 mM EGTA, 0.1 mM EDTA). Bound proteins were eluted with a 45 ml of linear gradient of 0-1 M KCl in HB buffer. Fractions containing the proteins were combined and concentrated on amicon spin filters with a 50 kDa cutoff after addition of 0.1 mM ATP and 10% glycerol. Concentrated proteins were frozen in LiN2 and stored at −80°C.

KLC1, SKIP-1-310, JIP1 were purified via size exclusion chromatography using Superose 6 10/300 increase GL (GE Healthcare) column equilibrated in lysis buffer. KLC1-CC, KLC1-TPR, KLC1, Nesprin-4 and Nesprin-4-WD/AA were purified via size exclusion chromatography using Superdex 200 10/300 GL (GE Healthcare) column equilibrated in lysis buffer. Fractions containing the proteins were combined and concentrated on amicon spin filters with a 50 kDa cutoff. Concentrated proteins were frozen in LiN2 and stored at −80°C.

KIF5A, KIFB, KIF5C, KIF5B-Δhinge, KIF5B-IAK/AAA, KIF5B-E178A, KIF5A-KLC1, KIF5B-KLC1, KIF5B-KLC1 were purified via size exclusion chromatography using BioSep SEC-s4000 (Phenomenex) particle size 5 μm, pore size 500Å, 7.8 mm ID x 600 mm column equilibrated in GF150 buffer (25 mM HEPES-KOH, 150 mM KCl, 2 mM MgCl2). Fractions containing the proteins were combined and concentrated on amicon spin filters with a 50 kDa cutoff after addition of 0.1 mM ATP and 10% glycerol. Concentrated proteins were frozen in LiN2 and stored at −80°C. sfGFP-MAP7 protein was purified from insect cells as previously described (Monroy et al., 2018).

### SEC-MALS (size exclusion chromatography coupled to multiangle light scattering)

The purified proteins were analyzed using BioSep SEC-s4000 (Phenomenex) particle size 5 μm, pore size 500Å, 7.8 mm ID x 300 mm column equilibrated in GF150 buffer (25 mM HEPES-KOH, 150 mM KCl, 2 mM MgCl2) plumbed into an HPLC (Agilent 1100). Molecular masses were analyzed by an inline SEC-MALs system (Wyatt Technology) which included a miniDAWN TREOS to measure light scattering and an Optilab T-rEX to measure refractive index. Molar mass was calculated using ASTRA v. 6 software (Wyatt Technology).

### Microtubule assembly

Porcine brain tubulin was isolated using the high-molarity PIPES procedure and then labelled with biotin NHS ester, Dylight 650 NHS ester or Dylight-405 NHS ester as described previously (http://mitchison.hms.harvard.edu/files/mitchisonlab/files/labeling_tubulin_and_quantifying_labeling_stoichiometry.pdf). Pig brains were obtained from a local abattoir and used within ~4 h after death. Microtubules were prepared by mixing 60 μM unlabeled tubulin, 3 μM biotin-labeled tubulin and 3 μM Dylight-405-labeled tubulin for single molecule assays and 60 μM unlabeled tubulin and 3 μM Dylight-650-labeled tubulin for gliding assays. Mixed tubulin were incubated at 37°C with 10 mM GTP for 15 min. Then 20 μM taxol was added to stabilize the polymerized microtubule and additional incubation was performed at 37°C for 30 min. Microtubules were pelleted by centrifugation at 20,000 x g for 10 min over a 25% sucrose cushion and the pellet was resuspended in 50 μl BRB80 (80 mM PIPES-KOH pH 6.8, 1 mM MgCl2 and 1 mM EGTA) containing 10 μM taxol.

### TIRF assays

Glass chambers were prepared by acid washing as previously described (Tan et al., 2017). Dylight-405/biotin labeled microtubules were flowed into streptavidin adsorbed flow chambers and allowed to adhere for 5–10 min. Unbound microtubules were washed away using assay buffer (90 mM HEPES-KOH pH 7.6, 50 mM KCH_3_COO, 2 mM Mg(CH_3_COO)_2_, 1 mM EGTA, 10 % glycerol, 0.1 mg/ml biotin–BSA, 0.2 mg/ml K-casein, 0.5 % Pluronic F127, 2 mM ATP, and an oxygen scavenging system composed of PCA/PCD/Trolox). Purified motor protein was diluted to indicated concentrations in the assay buffer. Then, the solution was flowed into the glass chamber. Sequential images of 488 and/or 561 channel were taken at the frame rate of 1 fps (Fig. 6A, 6B and 6F), 2 fps (Fig. 2E, 3E and 5C) or 10 fps (Fig. 1C, 2I, 3H, 4C and 5D), respectively. Assays using AMP-PNP were performed using 2 mM AMP-PNP instead of ATP in the above. Prior to taking images, chambers were incubated at room temperature for 10 min to allow proteins time to react with AMP-PNP. For gliding assay, motor proteins were diluted in lysis buffer at 0.5 μM and first flowed into empty chambers. After incubating at room temperature for 10 min, residual proteins were washed away with assay buffer. Then 650-labeled microtubules diluted in assay buffer were flowed in. Microtubule channel were taken at 2 fps. Images were acquired using a Micromanager (Edelstein et al., 2010) software-controlled Nikon TE microscope (1.49 NA, 100× objective) equipped with a TIRF illuminator and Andor iXon CCD EM camera. Statistical tests were performed in GraphPad PRISM 8.

### Pull-down assays

Pull-down assays were performed with purified Nesprin-4, Nesprin-4-WD/AA. Rat brain was homogenized in equal weight:volume of buffer (50 mM Hepes, 50 mM Pipes pH 7.0, 1 mM EDTA, 2 mM MgSO4) using a dounce homogenizer and flash frozen in LiN2 and stored at −80°C. The frozen lysate was thawed on ice and supplemented with 2 mM DTT, 2 mM PMSF and 0.1% Nonidet P-40, followed by centrifugation at 270,000 x g for 10 min at 4°C. The supernatant was mixed with 300 nM of the StrepII-tagged proteins and incubated for 30 min on ice and the protein complexes were recovered by Strep agarose beads. The beads were washed with wash buffer (90 mM HEPES-KOH pH 7.6, 50 mM KCH_3_COO, 2 mM Mg(CH_3_COO)_2_, 1 mM EGTA, 10 % glycerol, 0.1% NP-40) for 5 times and eluted with 3 mM d-desthiobiotin in wash buffer. The eluates were analyzed by SDS-PAGE and western blotting.

### Analysis of protein sequences

Protein sequences were aligned by Jalview 2 and Clustal W. Identities between protein sequences were calculated with the SMS2 tool (https://www.bioinformatics.org/sms2/ident_sim.html).

## Author Contributions and Notes

RJM, KC, and SN designed research. RJM and KMOM secured research funding, KO performed research and analyzed data. RJM and KO wrote the paper. All authors edited the paper. The authors declare no conflict of interest.

## Acknowledgments

The authors thank all the members of the MOM lab for their continual input and feedback on this project. We thank Dan Starr (UC Davis) for generously providing the cDNA for human nesprin-4. The work was supported by grants from NIGMS GM124889 (to RJM) and GM133688 (to KMOM.), The Japan Society for the Promotion of Science 20H03247, 19H04738, and 16H06536 (to S.N.). The Osamu Hayaishi Memorial Scholarship for Study Abroad (to K.C.), and a JSPS Overseas Research Fellowship (to K.C.).

## Supplementary Figure S1

**Supplementary Figure S1.**
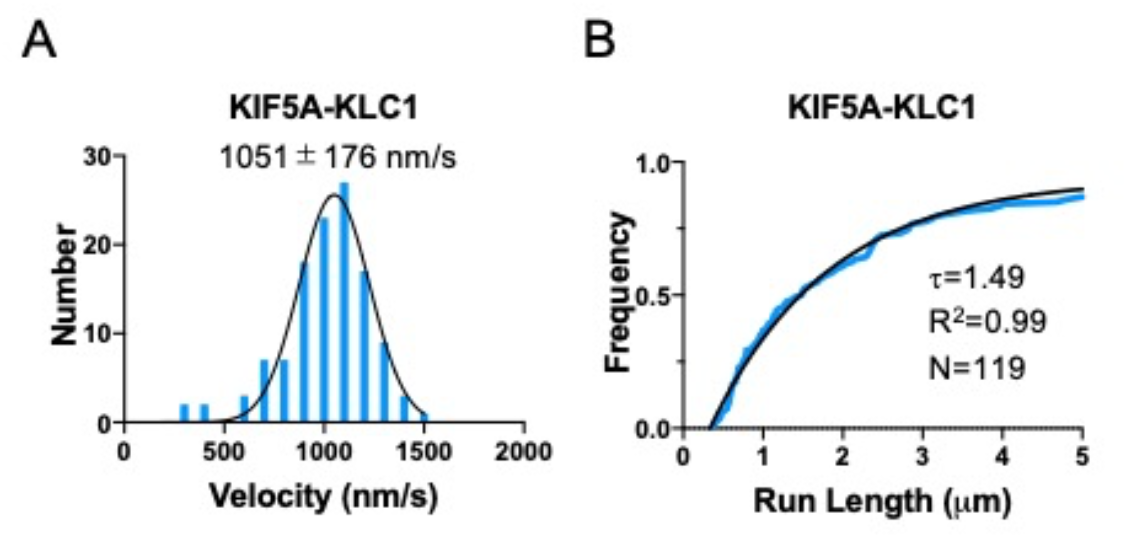
**(A)** Velocity histogram with Gaussian fit of processive KIF5A-KLC1 motors. Mean ± SD is 1051 ± 176 nm/s. **(B)** Run lengths plotted by 1-cumulative frequency distribution with one-phase exponential decay fit. Decay constant (τ, run length) is 1.49 μm. R^2^ = 0.99. N = 2. n = 119 molecules.

## Supplementary Figure S2

**Supplementary Figure S2.**
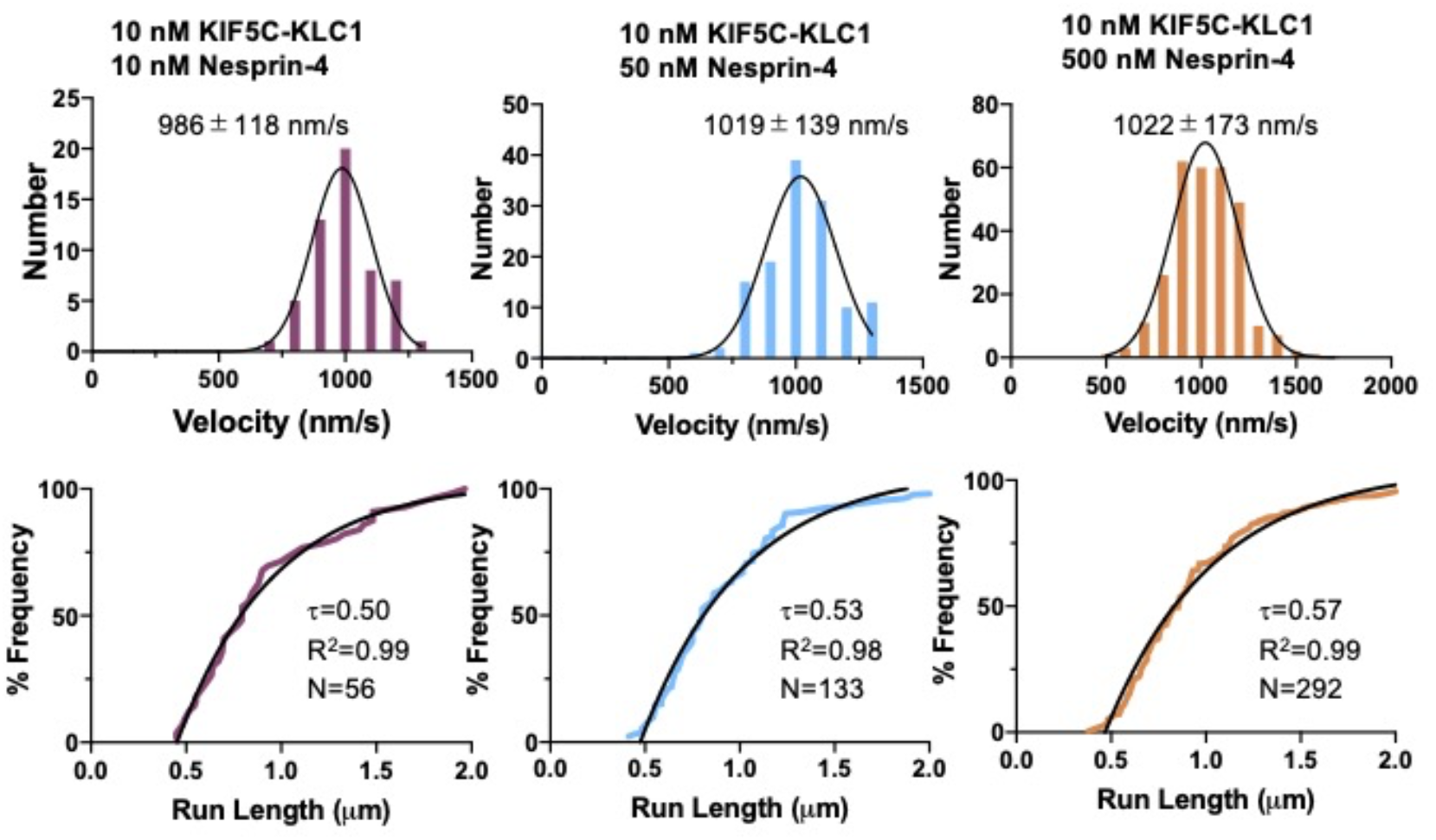
**(A)** Velocity histograms with Gaussian fits of KIF5C-KLC1 tetramers with or without nesprin-4. Means ± SD: 986 ± 118 μm/s (10 nM KIF5C-KLC1 + 10 nM Nesprin-4), 1019 ± 139 μm/s (10 nM KIF5C-KLC1 + 50 nM nesprin-4) and 1022 ± 173 μm/s (10 nM KIF5C-KLC1 + 500 nM Nesprin-4). N = 2. n = 56, 133 and 292 molecules, respectively. **(B)** Run lengths plotted by 1-cumulative frequency distribution with one-phase exponential decay fit. Decay constant (τ, run length) and R^2^ value of the fits are shown.

